# pVHL regulates protein stability of TCF/LEF transcription factor family via ubiquitin-independent proteasomal degradation

**DOI:** 10.1101/2021.10.14.464355

**Authors:** Caixia Wang, Xiaozhi Rong, Haifeng Zhang, Bo Wang, Yan Bai, Yunzhang Liu, Ling Lu, Yun Li, Yonghua Sun, Chengtian Zhao, Jianfeng Zhou

## Abstract

The Wnt/β-catenin signaling pathway plays key roles in development and adult tissue homeostasis by controlling cell proliferation and cell fate decisions. In this pathway, transcription factors TCF/LEFs are the key components to repress target gene expression by recruiting co-repressors or to activate target gene expression by recruiting β-catenin when the Wnt signals are absent or present, respectively. While progress has been made in our understanding of Wnt signaling regulation, the underlying mechanism that regulates the protein stability of the TCF/LEF family is far less clear. Here, we show that von Hippel-Lindau protein (pVHL), which is the substrate recognition component in an E3 ubiquitin ligase complex, controls TCF/LEF protein stability. Unexpectedly, pVHL directly binds to TCF/LEFs and promotes their proteasomal degradation independent of E3 ubiquitin ligase activity. Knockout of *vhl* in zebrafish embryos leads to a reduction of dorsal habenular neurons and this effect is upstream of dorsal habenular neurons phenotype in *tcf7l2*-null mutants. Our study uncovers a previously unknown mechanism for the protein stability regulation of the TCF/LEF transcription factors and demonstrates that pVHL contains a 26S proteasome binding domain that drives ubiquitin-independent proteasomal degradation. These findings provide new insights into the ubiquitin-independent actions of pVHL and uncover novel mechanistical regulation of Wnt/β-catenin signaling.

## Introduction

The Wnt/β-catenin signaling pathway is an evolutionarily conserved signal transduction cascade that controls numerous developmental processes and plays crucial roles in the regulation of diverse processes, including stem cell renewal, cell proliferation, and cell differentiation during adult tissue homeostasis in multicellular animals(Clevers et al., 2014; Clevers and Nusse, 2012; MacDonald et al., 2009; Nusse and Clevers, 2017; Steinhart and Angers, 2018). Dysregulation of the Wnt/β-catenin signaling cascade is often associated with various kinds of human diseases, including many cancers and hereditary diseases(Anastas and Moon, 2013; Clevers and Nusse, 2012; MacDonald *et al*., 2009; Nusse and Clevers, 2017). In the absence of Wnt ligands, cytosolic β-catenin interacts with a destruction complex consisting of APC, GSK3, CK1, AXIN1, and β-TrCP, which leads to the phosphorylation of N-terminal Ser/Thr residues of β-catenin by CK1 and GSK3. Consequently, β-catenin is ubiquitylated and undergoes proteasome-mediated degradation to maintain the cytoplasmic β-catenin at low levels(Clevers and Nusse, 2012; MacDonald *et al*., 2009; Stamos and Weis, 2013). Once Wnt ligands bind to the Frizzled family transmembrane receptors and LRP5/6 coreceptors, the disaggregation of the destruction complex is triggered. Consequently, β-catenin is non-phosphorylated and stabilized, which allows it to accumulate in the cytoplasm and translocate into the nucleus. In the nucleus, DNA-bound TCF/LEF transcription factors act as transcriptional repressors by interacting with Groucho proteins, while they can transiently convert into transcriptional activators upon β-catenin engagement(Clevers and Nusse, 2012; MacDonald *et al*., 2009; Nusse and Clevers, 2017). Thus, the ultimate outcome of the Wnt signal is determined by β-catenin and the TCF/LEFs.

All TCF/LEF family members are high-mobility group DNA-binding proteins with multiple domains for protein interaction and regulation(Clevers and Nusse, 2012; MacDonald *et al*., 2009). The TCF/LEFs possess a highly conserved high-mobility group DNA-binding domain (HMG DBD), which consists of an HMG box and a nuclear localization signal and can recognize and bind specific DNA sequences(Cadigan and Waterman, 2012; Doumpas et al., 2019). To date, four TCF protein family members have been identified in vertebrate genomes. Currently available evidence suggests that the relative amounts of β-catenin and TCF in the nucleus influence the Wnt signaling output(Goentoro and Kirschner, 2009; Phillips and Kimble, 2009). Hence, nuclear TCF/LEF concentrations may be dynamically controlled as precisely as that of β-catenin(Cadigan and Waterman, 2012). Previous reports have indicated that certain TCF/LEFs are implicated in the ubiquitin-proteasome pathway(Ishitani et al., 2005; Shy et al., 2013; Yamada et al., 2006). For example, NARF, a nemo-like kinase (NLK)-associated RING finger protein, is an E3 ubiquitin ligase that regulates TCF7L2 and LEF1 ubiquitylation and degradation(Yamada *et al*., 2006). Compared with the well understood protein stability regulation of β-catenin as an effector of the Wnt signaling pathway, however, the stability of the TCF/LEF proteins is far less clear.

Wnt/β-catenin signaling acts as a key regulator in the neurogenesis of the habenula(Beretta et al., 2013; Carl et al., 2007; Husken and Carl, 2013; Husken et al., 2014). In zebrafish, habenular neuron types are grossly divided into dorsal and ventral habenular neurons. The dorsal habenular neuronal clusters (dHb) consist of lateral (dHbl) and medial (dHbm) subnuclei. The development of dHb is asymmetrical with a left-right difference. The dHbl subnuclei are larger on the left, whereas the dHbm subnuclei are larger on the right(Concha and Wilson, 2001). Distinct temporal regulation of Wnt/β-catenin signaling is pivotal for the diversity of cell fate during the asymmetric development of dorsal habenula. Premature activation of Wnt/β-catenin signaling between 24 and 26 hpf can lead to delayed differentiation of dHb neurons, resulting in significant reduction in early born dHbl neurons and a mild reduction in late born dHbm neurons(Guglielmi et al., 2020). In Wnt-overactivated zebrafish *axin1* mutants, dHbl charactered double-right-sided, which was opposite to dHbl with double-left-sided character in *tcf7l2*-null mutant(Carl *et al*., 2007; Husken *et al*., 2014). Additionally, inhibition of Wnt/β-catenin signaling between 34 and 36 hpf leads to an induction of dHbl neurons on the right side(Husken *et al*., 2014).

The von Hippel-Lindau protein (pVHL) is the target protein recognition subunit of an E3 ubiquitin ligase complex (VBC, including pVHL and elongins B and C)(Kaelin, 2007). Various natural mutations with inactivation of the *VHL* gene, a tumor suppressor gene, have been reported in most cases of hereditary von Hippel-Lindau disease and sporadic clear-cell renal carcinomas (ccRCCs)(Gossage et al., 2015; Kaelin, 2007). pVHL targets prolyl-hydroxylated proteins(Guo et al., 2016; Ivan et al., 2001; Jaakkola et al., 2001; Zhang et al., 2018). Hypoxia-inducible factors (HIF-α), which are prolyl-hydroxylated by EGLN 1/2/3 family proteins under normal oxygen tension, are the well-documented canonical targets of pVHL. Hydroxylated HIF-α is bound and recognized by pVHL for ubiquitination-mediated proteasomal degradation(Ivan *et al*., 2001; Jaakkola *et al*., 2001). An earlier study revealed that pVHL binds to DVL and ubiquitinates it, ultimately promoting DVL degradation via the autophagy-lysosome pathway(Gao et al., 2010). In addition, pVHL facilitates β-catenin degradation by stabilizing Jade-1(Berndt et al., 2009; Chitalia et al., 2008). Despite the multiple pVHL functions in the process of Wnt signaling regulation, it is unknown whether pVHL degrades proteins independently of the ubiquitin-proteasome and autophagy-lysosome pathways and what the underlying mechanism is.

We previously reported that either overexpression or knockdown of VBP1, a pVHL binding protein, increased the association between pVHL and TCF/LEFs and attenuated Wnt/β-catenin signaling via the promotion of the proteasomal degradation of TCF/LEFs(Zhang et al., 2020). In this process, pVHL is required in the effects of VBP1 on the stability of TCF/LEFs. Intriguingly, we observed that VBP1 enhances the ubiquitylation of HIF-1α but fails to induce the ubiquitylation of Tcf7l2. This suggested that pVHL likely promotes the proteasomal degradation of TCF/LEFs with a previously unreported mechanism. In this study, we aim to elucidate the underlying mechanisms of pVHL action on TCF/LEFs as well as the physiological role of this action in a model organism.

## Results

### pVHL inhibits Wnt/β-catenin signaling and promotes TCF/LEFs protein degradation

pVHL has been suggested to 1) facilitate the ubiquitylation and autophagy-mediated degradation of DVL2 and 2) downregulate β-catenin through the mediator Jade-1 to reduce predominantly phospho-β-catenin and inhibit Wnt/β-catenin signaling(Chitalia *et al*., 2008; Gao *et al*., 2010). In addition, our previous study reported that VBP1 regulates TCF/LEF protein stability via pVHL(Zhang *et al*., 2020). To investigate the underlying mechanism of pVHL regulation of TCF/LEF protein stability, we examined the effects of pVHL on Wnt-induced TCF/LEF-dependent transcriptional activity. A TOPFlash reporter plasmid, which contained Wnt-responsive TCF/LEF binding sites, was transfected into HCT116 cells, and then the transcriptional activity was measured. We observed that pVHL decreased expression of the TOPFlash reporter in HCT116 cells in a dose-dependent manner (Figure 1A). This observation was surprising since HCT116 cells are β-catenin constitutively activated by a Ser45 deletion in one of the β-catenin alleles(Li et al., 2012). We then tested whether pVHL modulates Wnt/β-catenin signaling at the TCF/LEF level. To this end, pVHL and constitutively active Tcf7l1 (VP16-Tcf7l1ΔN, a β-catenin-independent VP16-Tcf7l1 fusion protein lacking the β-catenin-binding site) were co-transfected into HEK293T cells, and Wnt reporter activity was monitored. In agreement with the inhibitory effect of VBP1 on VP16-Tcf7l1ΔN-induced Wnt reporter activity previously reported(Zhang *et al*., 2020), pVHL also inhibited VP16-Tcf7l1ΔN-induced Wnt reporter activity in a dose-dependent manner (Figure 1B).

**Figure 1.**
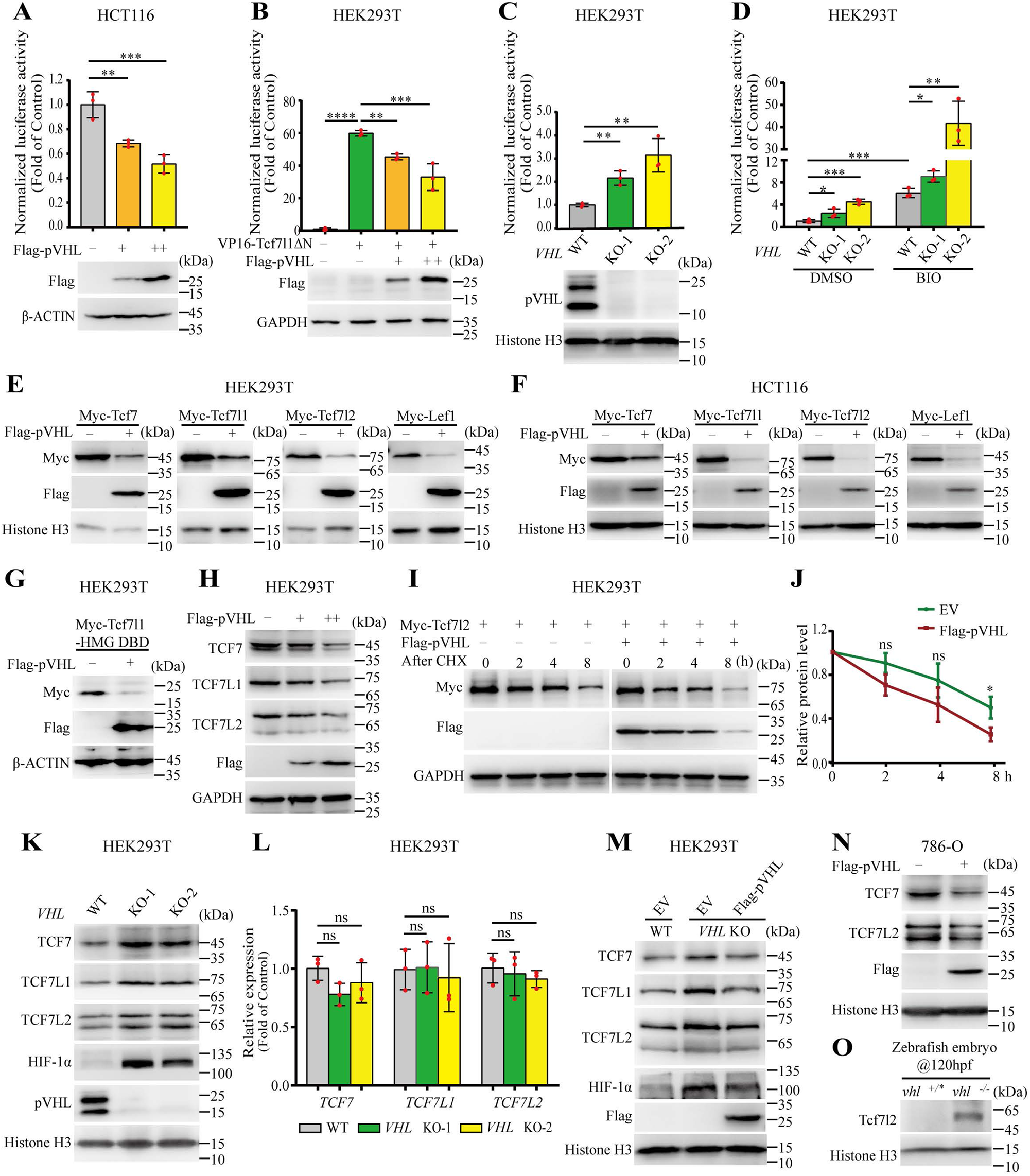
pVHL inhibits Wnt/β-catenin signaling and stabilizes TCF/LEF protein. (A) TOPFlash luciferase assays in HCT116 cells with increasing pVHL overexpression. Values are mean ± S.D. (n=3). One-way ANOVA analysis with Dunnett’s multiple comparisons test, \*\**p* < 0.01; \*\*\**p* < 0.001. (B) TOPFlash assays in VP16-Tcf7l1ΔN-treated HEK293T cells with increasing pVHL overexpression. Wnt/β-catenin signal was activated by transfection with 50 ng VP16-Tcf7l1ΔN plasmid DNA. Expression of Flag-pVHL was confirmed by western blotting. Values are mean ± S.D. (n=3). One-way ANOVA analysis with Dunnett’s multiple comparisons test, \*\**p* < 0.01; \*\*\**p* < 0.001; \*\*\*\**p* < 0.0001. (C) TOPFlash luciferase assays in *VHL*-knockout HEK293T cells. pVHL protein levels were confirmed by western blotting. Values are mean ± S.D. (n=3). Unpaired *t*-test, \*\**p* < 0.01. (D) TOPFlash luciferase assays in BIO-treated *VHL*-knockout HEK293T cells. TOPFlash plasmid was cotransfected with *Renilla* plasmid into control or *VHL*-knockout cells. Wnt/β-catenin activity was induced by BIO (1 μM). Values are mean ± S.D. (n=3). Unpaired *t*-test, \**p* < 0.05; \*\**p* < 0.01; \*\*\**p* < 0.001. (E, F) Exogenous Tcf/Lef protein levels in control or pVHL-overexpressing HEK293T and HCT116 cells. (G) Tcf7l1-HMG DBD protein level in HEK293T cells with Flag-pVHL overexpression. Endogenous TCF protein levels in HEK293T cells with increasing pVHL overexpression. Flag-pVHL promotes Myc-Tcf7l2 degradation in HEK293T cells. HEK293T cells were transfected with Myc-Tcf7l2, co-transfected with empty vector or Flag-pVHL, after 16 h, treated with cycloheximide (CHX; 100 μg/mL), and harvested at indicated time points. (J) Quantification of (I). Myc-Tcf7l2 protein level normalized to GAPDH. Values are mean ± S.D. (n=3). Two-way ANOVA analysis with Bonferroni’s multiple comparisons test, ns, not significant; \**p* < 0.05. (K) The protein levels of TCFs in control and *VHL*-Knockout cells. The expression level of HIF-1α was used as a positive control. (L) The transcriptional levels of *TCF*s in control and *VHL*-knockout cells were analyzed by qRT-PCR. Values are mean ± S.D. (n=3). Unpaired *t*-test, ns, not significant. (M) Introduction of Flag-pVHL into *VHL*-knockout HEK293T cells downregulated TCFs protein levels. (N) Reintroduction of Flag-pVHL downregulated TCF7 and TCF7L2 in 786-O cells. (O) *vhl*-deficiency upregulated Tcf7l2 protein in zebrafish embryos. Sibling and mutant embryos were harvested at 120 hpf. Protein samples of 4 zebrafish embryos were added in each well.

To further confirm these results, *VHL*-knockout HEK293T cells were generated using a CRISPR/Cas9-mediated gene editing approach (Figure 1-figure supplement 1A, B). Two pVHL protein isoforms, a long form and a short form, have been previously reported(Kaelin, 2007). The generated *VHL*-knockout lines have a premature termination codon at exon 1, which causes both isoforms to be depleted (Figure 1C). We measured Wnt luciferase reporter activity after depletion of pVHL. As expected, knockout of pVHL enhanced basal Wnt reporter activity (Figure 1C). HEK293T cells were regarded as a Wnt-off cell line. Addition of a GSK3 inhibitor, 6-bromoindirubin-3’-oxime (BIO), in HEK293T cells could induce Wnt reporter activity (Figure 1D). To further examine the effect of *VHL* depletion on Wnt activity, *VHL-*depleted HEK293T cells were treated with BIO, and the Wnt reporter activity was measured. Knockout of pVHL further increased Wnt reporter activity in the background of BIO treatment (Figure 1D). Taken together, these results suggested that pVHL inhibits Wnt/β-catenin signaling.

pVHL is the substrate recognition component of an E3 ubiquitin ligase complex. Additionally, VBP1 regulates TCF/LEF protein degradation via pVHL. We speculate that pVHL may promote VP16-Tcf7l1ΔN protein degradation and prevent its ability to induce the Wnt reporter activity. To test this hypothesis, we co-transfected Flag-tagged pVHL with Myc-tagged Tcf7, Tcf7l1, Tcf7l2, and Lef1 into HEK293T and HCT116 cells, which were under Wnt-off and Wnt-on conditions, respectively. The overexpression of pVHL reduced the abundance of Tcf7, Tcf7l1, Tcf7l2, and Lef1 in both cell lines (Figure 1E and F). The HMG DBD (DNA binding domain) of TCF/LEFs is evolutionarily conserved and nearly identical from invertebrate to vertebrate(Cadigan and Waterman, 2012). Given that HMG DBD is the most highly conserved domain in TCF/LEFs and that pVHL can reduce the abundance of four TCF/LEFs, we assume that pVHL may also downregulate the HMG DBD protein. To prove this hypothesis, pVHL and Myc-tagged Tcf7l1-HMG DBD were co-transfected into HEK293T cells. The Tcf7l1-HMG DBD protein levels were dramatically reduced by pVHL overexpression (Figure 1G). Consistent with these results, pVHL decreased the abundance of endogenous TCF7, TCF7L1, and TCF7L2 proteins in the HEK293T cells in a dose-dependent manner (Figure 1H). Therefore, pVHL negatively regulates Wnt/β-catenin activity and the TCF/LEF protein level.

To determine whether pVHL promotes TCF/LEF protein degradation, we performed a time-course treatment assay with cycloheximide (CHX), an inhibitor of protein synthesis. When Flag-tagged pVHL was co-transfected with Myc-tagged Tcf7l2, the degradation of Myc-tagged Tcf7l2 was accelerated (Figure 1I and J). To verify the specificity of pVHL, we measured protein levels of TCF7, TCF7L1, and TCF7L2 in *VHL-*knockout HEK293T cells. Knockout of pVHL robustly increased HIF-1α protein levels. Likewise, knockout of pVHL markedly increased TCF7, TCF7L1, and TCF7L2 protein levels (Figure 1K). Quantitative real-time RT-PCR analysis showed that knockout of pVHL did not alter the mRNA levels of *TCF7, TCF7L1*, or *TCF7L2* (Figure 1L). Moreover, reintroduction of pVHL into *VHL*-knockout HEK293T cells neutralized this effect, as the TCF7, TCF7L1, TCF7L2, and HIF-1α protein levels were significantly reduced (Figure 1M). To validate this result, we further reintroduced pVHL into *VHL*-deficient ccRCC 786-O cells. The reintroduction of pVHL led to a reduction in endogenous TCF7 and TCF7L2 protein levels (Figure 1N). These results suggested that the *VHL* knockout is specific and that pVHL promotes TCF/LEF protein degradation *in vitro*.

We next investigated the effects of pVHL on TCF/LEF degradation *in vivo*. Zebrafish pVhl is an ortholog of the short human pVHL isoform(van Rooijen et al., 2009). We subjected zebrafish *vhl*-null mutant embryos to western blot to determine their Tcf7l2 protein levels. Indeed, knockout of pVhl in zebrafish caused accumulation of Tcf7l2 at the larval stage (120 hpf) (Figure 1O). Taken together, these results suggested that pVHL regulates TCF/LEFs abundance *in vitro* and *in vivo* and that the effects of pVHL/pVhl on TCF/LEFs are evolutionarily conserved.

### pVHL interacts with TCF/LEFs and promotes their proteasomal degradation

To examine whether TCF/LEFs and pVHL interact with each other, four Myc-tagged TCF/LEF members were expressed in HEK293T cells, and cell lysates were subjected to immunoprecipitation (IP) with anti-Myc antibody. Co-IP assay showed that endogenous pVHL interacts with all four TCF/LEF members (Figure 2A). In addition, co-IP assay also indicated that endogenous TCF7L2 retrieved endogenous pVHL in HEK293T cells (Figure 2B). Furthermore, purified glutathione-S-transferase (GST)-pVHL protein pulled down all four Myc-tagged TCF/LEFs *in vitro* (Figure 2C). Likewise, a direct protein-protein interaction between pVHL and TCF7L2 was also confirmed by a GST pulldown assay (Figure 2D). Collectively, these data revealed that TCF/LEFs and pVHL directly interact with each other.

**Figure 2.**
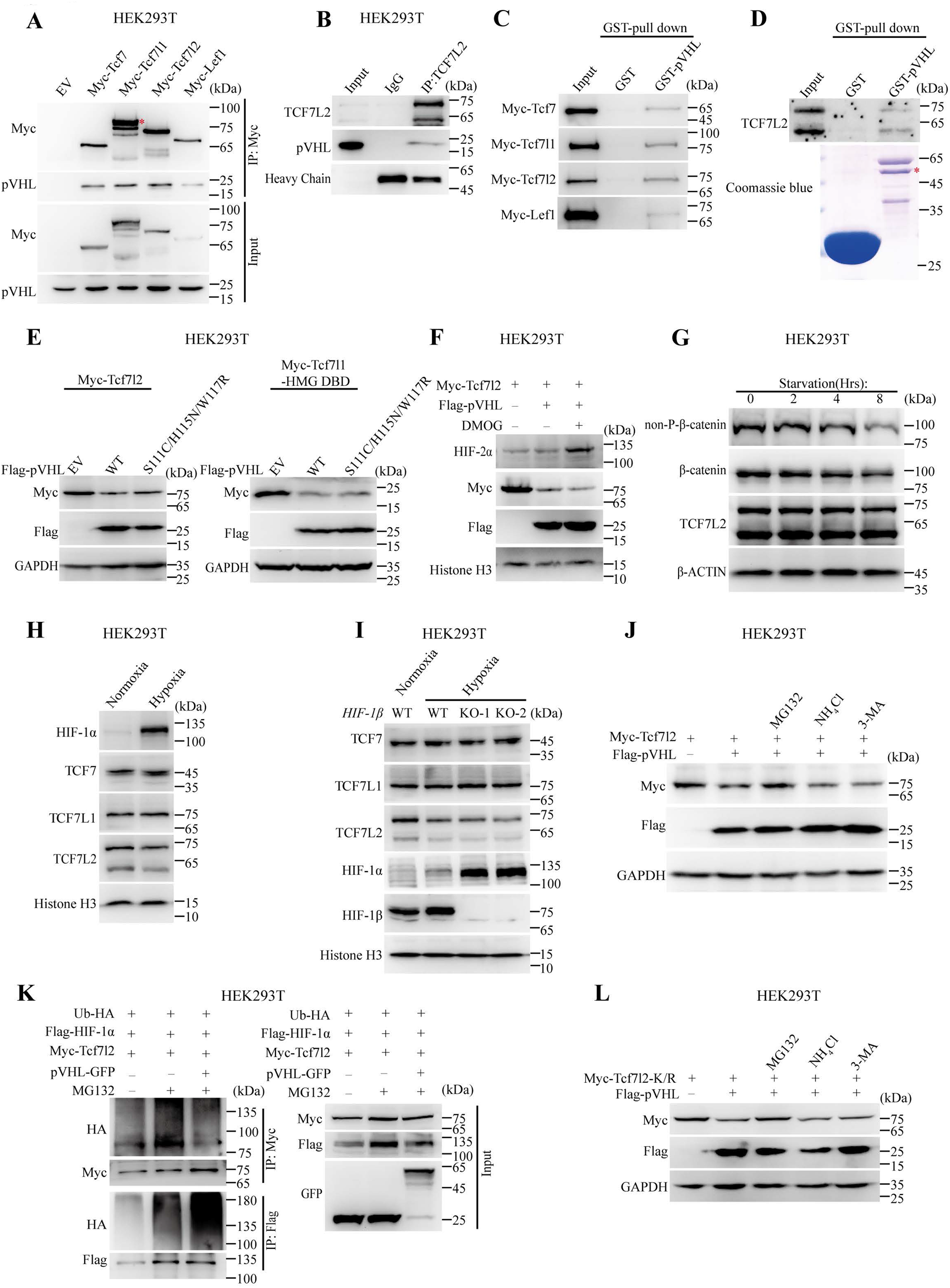
pVHL directly binds with TCF/LEFs and promotes their degradation by ubiquitin-independent proteasome pathway. (A) Detection of pVHL binding to TCF/LEFs in HEK293T cells by Co-IP. Red asterisk indicates the specific band. (B) Co-IP assay revealed the endogenous interaction between TCF7L2 and pVHL in HEK293T cells. IgG heavy chain was used as a negative control. (C, D) pVHL directly binds with TCF/LEFs. Purified GST or GST-pVHL proteins were incubated with extracts of HEK293T cells either transfected with Myc-Tcf/Lefs (C) or untransfected (D). Bound proteins were eluted and analyzed by western blot using indicated antibodies. (E) Tcf7l2 and Tcf7l1-HMG DBD protein levels in HEK293T cells with pVHL- or pVHL-S111C/H115N/W117R-overexpression. (F) pVHL promoted Tcf7l2 degradation in presence of the DMOG. Western blot analysis of WCL derived from HEK293T cells transfected with indicated plasmid DNA and either untreated or treated with 200 μM DMOG for 12 h. (G) Time-course for non-phospho-(active) β-catenin, β-catenin, and TCF7L2 protein levels in starved HEK293T cells. Western blot analysis of WCL derived from starved HEK293T cells at indicated time points. (H, I) Endogenous TCF7, TCF7L1, TCF7L2, and HIF-1α protein level under normoxia (21% O_2_) or hypoxia (1% O_2_) condition for 24 h in wild-type (H) and in *HIF-1β*-knockout HEK293T cells (I). (J) Changes in Tcf7l2 protein levels in pVHL-overexpressing HEK293T cells treated with indicated inhibitors. The transfected cells were either untreated or treated with 10 μM MG132, 25 mM NH_4_Cl, or 5 mM 3-MA for 8 h. (K) Effects of pVHL-overexpression on Tcf7l2 ubiquitination. Myc-Tcf7l2, Flag-HIF-1α, and HA-Ub were co-transfected with GFP-Vector or pVHL-GFP into HEK293T cells. After 48h, cells were treated with 10 μM MG132 for 8 h, and lysed for immunoprecipitation with anti-Myc and anti-Flag antibody. Immunoprecipitates were detected by anti-Myc, anti-Flag, or anti-HA antibody. WCL were analyzed with anti-Myc, anti-GFP, or anti-Flag antibody. (L) Changes in Tcf7l2-K/R protein levels in pVHL-overexpressing HEK293T cells treated with indicated inhibitors. Western blot analysis of WCL derived from HEK293T cells transfected with indicated plasmid DNA and either untreated or treated with 10 μM MG132, 25 mM NH_4_Cl, or 5 mM 3-MA for 8 h.

pVHL recognizes and binds to prolyl-hydroxylated substrates, such as prolyl-hydroxylated HIF-α, Akt, and ZHX2, in order to exert its function. Three residues (S111, H115, and W117) in the pVHL hydroxyl-proline binding pocket are critical for pVHL interaction with prolyl-hydroxylated substrates such as prolyl-hydroxylated HIF-1α and Akt(Guo *et al*., 2016). Mutating these residues in pVHL abolishes its binding activity to prolyl-hydroxylated HIF-1α and Akt(Guo *et al*., 2016). The above triple residue-mutated pVHL was therefore utilized to test whether it has comparable functionality with that of the wild-type pVHL. Like wild-type pVHL, the pVHL mutant also downregulated abundance of Tcf7L2 (Figure 2E, left panel). Similarly, this pVHL mutant decreased the Tcf7l1-HMG DBD protein level when it was co-expressed with Tcf7l1-HMG DBD in HEK293T cells (Figure 2E, right panel). We next applied a prolyl hydroxylase inhibitor, dimethyloxalylglycine (DMOG), to inhibit the activity of EGLN 1/2/3. It has been reported that DMOG treatment inhibits the binding between HIF-2α and pVHL and stabilizes HIF-2α(Guo *et al*., 2016). This finding was consistent with our result (Figure 2F). However, DMOG treatment did not reverse Myc-tagged Tcf7l2 protein downregulation induced by pVHL overexpression (Figure 2F). Therefore, TCF/LEF degradation by pVHL does not depend on prolyl hydroxylation of TCF/LEFs.

A previous report has indicated that chronic starvation-stimulated autophagy negatively regulates Wnt/β-catenin signaling(Gao *et al*., 2010). We examined the effects of starvation, an autophagy stimulus with nutrient deprivation medium, on the expression of endogenous TCF7L2 in HEK293T cells. Chronic starvation reduced the protein levels of both non-p-β-catenin and total β-catenin but not that of TCF7L2 (Figure 2G). This result suggested that TCF7L2 is not degraded by autophagy.

Hypoxic exposure increases the levels of β-catenin and Wnt target effectors LEF1 and TCF7 by stabilizing HIFs in mouse embryonic stem cells(Mazumdar et al., 2010). However, the levels of β-catenin and TCF7L2 show no such effect in colorectal tumor cells(Kaidi et al., 2007). To exclude the effect of increased HIFs on TCF/LEF in *VHL*-knockout HEK293T cells, we examined the protein levels of TCFs with enhanced HIF-1α expression upon hypoxia treatment. Hypoxia treatment robustly increased the protein level of HIF-1α, while the protein levels of TCF7, TCF7L1, and TCF7L2 were not increased (Figure 2H). Dimerization of HIF-1α or HIF-2α with HIF-1β is mediated by their basic helix-loop-helix (bHLH) and PER-ARNT-SIM (PAS) domains, which are required for binding to hypoxia response elements (HREs) and HIF-dependent transcriptional activity(Wu et al., 2015). In this case, we generated *HIF1-β* (*ARNT*) knockout HEK293T cells, targeting exon 6 to disrupt its bHLH and PAS domains (Figure 2-figure supplement 1A, B). Compared with wild-type cells, the cells with absence of HIF-1β did not increase TCFs under hypoxia treatment (Figure 2I). Therefore, the level of HIF-α and HIF activity did not upregulate protein levels of TCFs.

To address the possible pathway of TCF/LEF degradation, we used specific small compound inhibitors, including MG132 (proteasomal inhibitor), NH_4_Cl (lysosomal proteolysis inhibitor), and 3-MA (autophagy inhibitor), to block the major protein degradation pathway. Addition of MG132 but not of NH_4_Cl or 3-MA blocked pVHL-mediated TCF/LEF degradation (Figure 2J). Thus, pVHL likely promotes TCF/LEF degradation via the proteasomal pathway. To verify the ubiquitylation of TCF/LEF, we performed *in vivo* ubiquitination assays to examine the effects of pVHL on Tcf7l2 ubiquitination. Myc-Tcf7l2, Flag-HIF-1α, and pVHL-GFP were co-transfected into HEK293T cells with HA-tagged ubiquitin. Similar to the effect of VBP1, pVHL overexpression increased the polyubiquitination level of HIF-1α while decreased that of Tcf7l2 (Figure 2K). Hence, pVHL did not function as a typical E3 ubiquitin ligase, which usually catalyzes its targets at its lysine (K) residue(s) to form polyubiquitin chains, to downregulate TCF/LEF in a manner similar to that for HIF-1α.

We further analyzed evolutionarily conserved lysine residues in several vertebrate TCF/LEF proteins and found approximately 19 conserved lysine residues in total (Figure 2-figure supplement 2). We mutated all of them into arginine residues (R) in a Myc-tagged *Xenopus* Tcf7l2 background (hereafter, Tcf7l2-K/R). We then determined whether pVHL could downregulate the Tcf7l2-K/R mutant. As expected, pVHL reduced the protein level of the Tcf7l2-K/R mutant. Moreover, addition of MG132 but not of NH_4_Cl or 3-MA blocked pVHL-mediated Tcf7l2-K/R mutant degradation (Figure 2L). Taken together, the above results suggested that pVHL promotes TCF/LEF proteasomal degradation independently of ubiquitin function.

To validate that pVHL-mediated TCF/LEF degradation does not rely on E3 ubiquitin ligase activity, we tested the effects on TCF/LEFs by naturally occurring and cancer-associated pVHL point mutants L158P and R167W and the truncated mutant pVHL (1-157). All three greatly reduced or diminished elongin B/C binding capability and abolished E3 ligase activity(Duan et al., 1995; Iwai et al., 1999; Ohh et al., 1998). When these mutants were co-expressed with Myc-tagged Tcf7l2 in HEK293T cells, they exhibited the same effects on the abundance of Tcf7l2 protein and VP16-Tcf7l1ΔN-induced Wnt reporter activity as wild-type pVHL (Figure 3A and B). Similar effects were observed when each mutant was co-expressed with Tcf7l1-HMG DBD (Figure 3C). We also examined whether these pVHL mutants without E3 ligase activity could downregulate the Tcf7l2-K/R mutant. Like wild-type pVHL, they all reduced the Tcf7l2-K/R mutant protein levels (Figure 3D). Moreover, we performed a CHX treatment time-course to test whether pVHL (1-157) overexpression accelerated Myc-tagged Tcf7l2 protein turnover. As wild-type pVHL, pVHL (1-157) also downregulated Tcf7l2 protein and shortened its half-life (Figure 3E and F).

**Figure 3.**
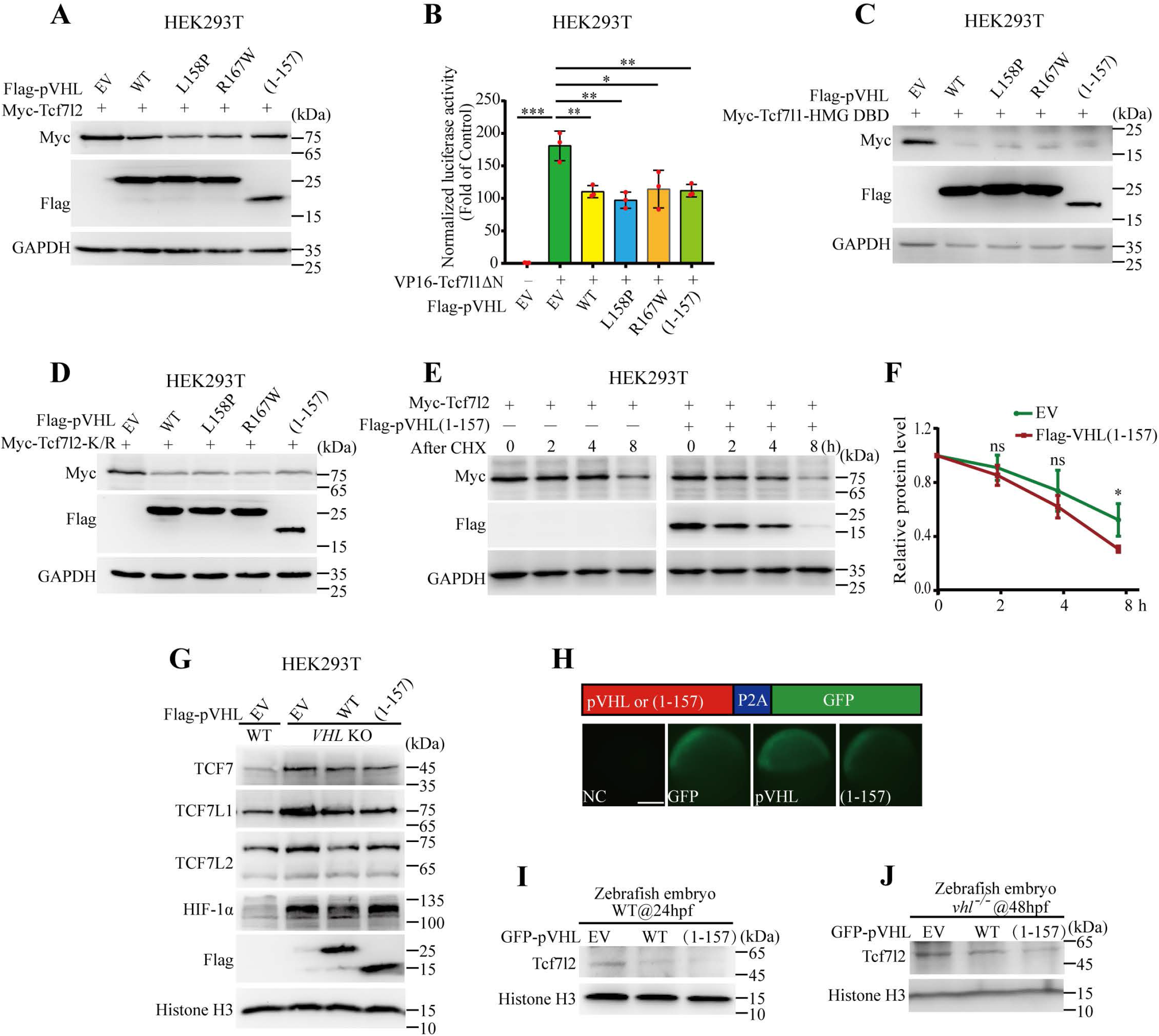
pVHL promotes TCF/LEF degradation in an E3 ubiquitin ligase-independent manner. (A) Tcf7l2 protein levels in HEK293T cells with overexpression of wild-type, site-mutated, or truncated pVHL. (B) TOPFlash reporter assays in VP16-Tcf7l1ΔN-transfected HEK293T cells with overexpression of wild-type, site-mutated, or truncated pVHL. Wnt/β-catenin signal was activated by transfection with 50 ng VP16-Tcf7l1ΔN. Values are mean ± S.D. (n=3). Unpaired *t*-test. \**p* < 0.05; \*\**p* < 0.01; \*\*\**p* < 0.001. (C) Tcf7l1-HMG DBD protein levels in HEK293T cells with overexpression of wild-type, site-mutated, or truncated pVHL. (D) Tcf7l2-K/R protein levels in HEK293T cells with overexpression of wild-type, site-mutated, or truncated pVHL. (E) pVHL truncation mutant pVHL(1-157) promotes Tcf7l2 degradation in HEK293T cells. HEK293T cells were transfected with Myc-Tcf7l2, along with either empty vector or Flag-pVHL(1-157), after 16 h, treated with cycloheximide (CHX; 100 μg/mL) and harvested at indicated time points. (F) Quantification of (E). Myc-Tcf7l2 protein level normalized to GAPDH. Values are mean ± S.D. (n=3). Two-way ANOVA analysis with Bonferroni’s multiple comparisons test. ns, not significant; \**p* < 0.05. (G) Overexpression of Flag-pVHL and Flag-pVHL(1-157) reduced TCF7, TCF7L1, and TCF7L2 protein levels in *VHL*-KO cells. HIF-1α was downregulated in *VHL*-KO after transfection with Flag-pVHL but not with Flag-pVHL(1-157). (H) GFP expression in zebrafish embryos injected with pVHL or pVHL(1-157) with self-cleaving P2A-GFP at 6 hpf. Embryos at 1-2 cell stage were injected with each indicated mRNA and raised to 6 hpf. Scale bar = 200 μm. (I) Overexpression of pVHL or pVHL(1-157) decreased Tcf7l2 protein level in wide-type zebrafish embryos at 24 hpf. Protein samples of 4 zebrafish embryos were added in each well. (J) Reintroduction of pVHL or pVHL(1-157) into *vhl*-null mutant zebrafish embryos reduced Tcf7l2 protein level at 48 hpf. Protein samples of 4 zebrafish embryos were added in each well.

We next introduced wild-type pVHL and the pVHL (1-157) mutant into *VHL-*knockout HEK293T cells and measured TCF/LEF protein levels. As with wild-type pVHL, introduction of pVHL (1-157) into *VHL* knockout HEK293T cells decreased TCF7, TCF7L1, and TCF7L2 protein accumulation by depleting *VHL*, while the HIF-1α protein level was reduced by wild-type pVHL rather than by the pVHL (1-157) mutant (Figure 3G). Therefore, pVHL (1-157) and wild-type pVHL had comparable effects on TCF/LEF downregulation. In addition, we used developing zebrafish embryos to determine the effects of human pVHL and pVHL (1-157) on the promotion of Tcf7l2 degradation *in vivo*. Thus, we generated *in vitro* transcribed *GFP, VHL-P2A-GFP*, or *VHL (1*-*157)-P2A-GFP* mRNA and injected them into zebrafish embryos. In the pVHL-P2A-GFP or pVHL (1-157)-P2A-GFP fusion proteins, GFP can be removed by the “self-cleaving” small peptide 2A and used as an indicator to confirm protein expression of pVHL and pVHL (1-157) in zebrafish embryos (Figure 3H). Like *GFP* mRNA-injected groups, the GFP signals in the *VHL-P2A-GFP* and *VHL (1*-*157)-P2A-GFP* mRNA-injected group could be observed at the shield stage, suggesting that pVHL and pVHL (1-157) proteins were successfully and highly expressed (Figure 3H). Western blot showed that the Tcf7l2 protein levels were dramatically reduced in wild-type zebrafish embryos injected with *VHL-P2A-GFP* or *VHL (1*-*157)-P2A-GFP* mRNA, indicating that the effect of pVHL/pVhl on Tcf7l2 is evolutionarily conserved *in vivo* (Figure 3I). In addition, introduction of human *VHL* mRNA also decreased Tcf7l2 protein levels in *vhl*-null mutant background (Figure 3J). The *VHL (1*-*157)* mRNA had a similar effect on Tcf7l2 in *vhl*-null mutant embryos (Figure 3J). Collectively, these data implied that pVHL does not function as an E3 ligase complex adaptor to promote TCF/LEF degradation.

### The pVHL substrate recognition domain is required to downregulate the TCF/LEF protein

To identify the functional pVHL domain(s) essential for promoting TCF/LEF degradation, various pVHL domain-deleted mutants were generated (Figure 4A). Using Co-IP assay, we mapped the domain(s) putatively responsible for the interaction between pVHL and TCF/LEFs. A region comprising amino acid (aa) residues 100-157 was required for its interaction with Tcf7l2 (Figure 4B). We then mapped the pVHL domain(s) required for promoting Tcf7l2 degradation. Deletion of aa 54-99 or aa 100-157 in pVHL diminished its downregulation effect on Tcf7l2 protein compared with that of wild-type pVHL (Figure 4C). We further investigated whether aa 54-99 or aa 100-157 of pVHL regulated Wnt reporter activity. Deletion of either region in pVHL diminished its inhibitory effect on VP16-Tcf7l1ΔN-induced Wnt reporter activity (Figure 4D). Deletion of aa 54-99 or aa 100-157 also abolished pVHL-mediated Tcf7l2-K/R mutant degradation ability (Figure 4E). Therefore, aa 54-99 or aa 100-157 in pVHL is necessary to promote Tcf/Lef degradation.

**Figure 4.**
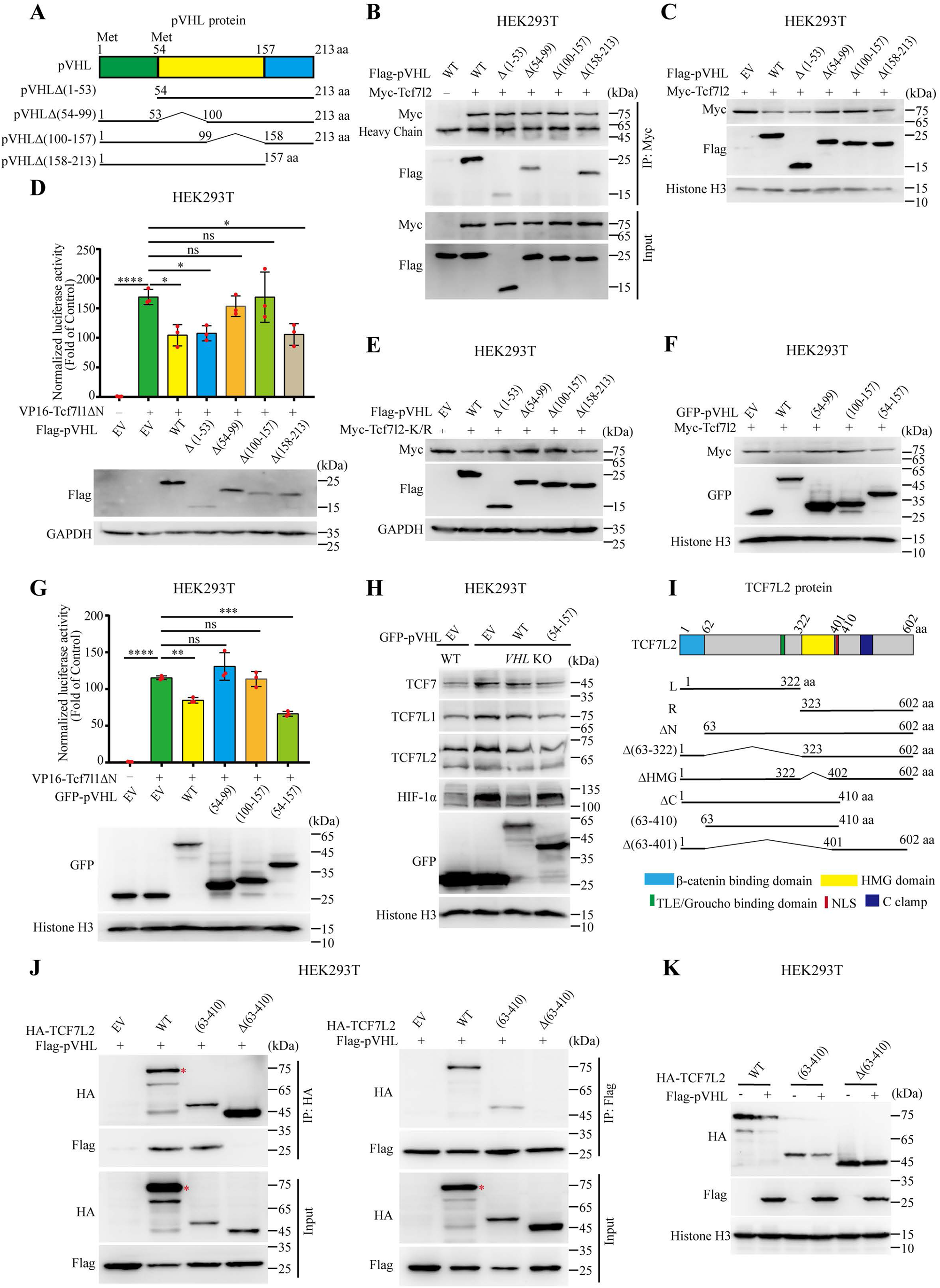
pVHL (54-157) is necessary and sufficient to promote TCF/LEF protein degradation. (A) Schematic representations of pVHL wild-type and truncated mutant proteins. (B) Mapping pVHL binding domain interacting with Tcf7l2 in transfected HEK293T cells by Co-IP assay. (C) Tcf7l2 protein levels in HEK293T cells with overexpression of indicated pVHL mutants. (D) TOPFlash reporter assays in HEK293T cells with coexpression of VP16-Tcf7l1ΔN and each indicated pVHL mutant. Wnt/β-catenin signal was activated by transfection with 50 ng VP16-Tcf7l1ΔN. Expression of the indicated Flag-tagged mutant pVHL was detected by an anti-Flag antibody. Values are mean ± S.D. (n=3). One-way ANOVA analysis with Dunnett’s multiple comparisons test. ns, not significant; \**p* < 0.05; \*\*\*\**p* < 0.0001. (E) Tcf7l2-K/R protein levels in HEK293T cells with overexpression of indicated pVHL mutants. Tcf7l2 protein levels in HEK293T cells with overexpression of indicated pVHL mutants. TOPFlash reporter assays in HEK293T cells with coexpression of VP16-Tcf7l1ΔN and indicated pVHL domains. Expression levels of indicated pVHL mutants detected by anti-GFP antibody. Values are mean ± S.D. (n=3). One-way ANOVA analysis with Dunnett’s multiple comparisons test. ns, not significant; ** *p* < 0.01; \*\*\**p* < 0.001; \*\*\*\**p* < 0.0001. (H) Introduction of pVHL and pVHL(54-157) into *VHL*-KO cells reduced TCF7, TCF7L1, and TCF7L2 protein levels. HIF-1α was downregulated in *VHL*-KO after transfection with pVHL-GFP but not with pVHL(54-157)-GFP. (I) Illustration of TCF7L2 wild-type and truncated mutant proteins. (J) Mapping TCF7L2 binding domain interacting with pVHL in transfected HEK293T cells by Co-IP assay. Red asterisk indicates the specific band. (K) The protein levels of TCF7L2 mutants in HEK293T cells with overexpressing pVHL.

To confirm whether aa 54-99 and aa 100-157 in pVHL suffice to promote TCF/LEF protein degradation, we generated GFP-tagged pVHL (54-99), pVHL (100-157), and pVHL (54-157) and evaluated their downregulation effects on TCF/LEF. Neither pVHL (54-99) nor pVHL (100-157) reduced Myc-tagged Tcf7l2 protein levels (Figure 4F). However, pVHL (54-157) was as effective at promoting TCF7L2 degradation as full-length pVHL (Figure 4F). pVHL (54-157) but not pVHL (54-99) or pVHL (100-157) inhibited VP16-Tcf7l1ΔN-induced TOPFlash reporter activity (Figure 4G). Additionally, pVHL (54-157) had the same capability as wild-type pVHL to reduce TCF/LEF protein levels in *VHL*-depleted HEK293T cells, while such effects were not observed on HIF-1α protein levels (Figure 4H). These results implied that pVHL (54-157) is necessary and sufficient to promote TCF/LEF protein degradation. In addition, these results also suggested that substrate recognition by pVHL as a component of E3 ubiquitin ligase is not required for TCF/LEF protein downregulation.

We next mapped the binding domain(s) of TCF/LEFs to pVHL. To define the domain(s) of TCF7L2 interacting with pVHL, a variety of domain-deleted mutants of TCF7L2 were generated based on conserved functional motifs (Figure 4I). To ensure the nuclear localization of each deletion mutant of TCF7L2, all of the mutants contained a nuclear localization signal (NLS). Co-IP analysis showed that deletion of aa 63-410, rather than deletion of other regions of TCF7L2, led to loss of the binding capability of TCF7L2 to pVHL (Figure 4J and Figure 4-figure supplement 1A-B). These results suggested that aa 63-410 of TCF7L2, containing both a TLE/Groucho binding domain and an HMG domain, is required for interaction with pVHL. Consistently, pVHL had little effect on the protein levels of the TCF7L2Δ(63-410) mutant, whereas overexpression of pVHL led to marked reduction in protein levels of other deletion mutants of TCF7L2 (Fig. 4K and Figure 4-figure supplement 1C). Immunostaining results confirmed that both TCF7L2(63-410) and TCF7L2Δ(63-410) mutants localized in the nucleus (Figure 4-figure supplement 1D). Altogether, we concluded that aa 63-410 of TCF7L2 is crucial for pVHL binding.

### pVHL directly interacts with the 26S proteasome

Several proteins, such as Parkin and Rad23, contain a ubiquitin-like domain (UBL), which is likely to interact and form a complex with RPN10 in the 19S regulatory subunit of the 26S proteasome and mediate substrate degradation(Hiyama et al., 1999; Sakata et al., 2003; Upadhya and Hegde, 2003). We hypothesized that pVHL has ubiquitin-independent proteasomal degradation activity to bridge TCF/LEF protein degradation. Therefore, we tested whether pVHL directly interacts with the 26S proteasome. An *in vitro* pulldown experiment on purified human 26S proteasome and recombinant GST-pVHL revealed that pVHL bound to the 19S regulatory subunit RPN10 and, therefore, directly interacted with the 26S proteasome (Figure 5A). Furthermore, we also performed a Co-IP assay in HEK293T cells with Flag-tagged pVHL to investigate whether pVHL and the 26S proteasome interact *in vivo*. As shown in Figure 5B, the endogenous 19S regulatory subunit RPN10 co-immunoprecipitated with pVHL. As deletion of either aa 54-99 or aa 100-157 disrupted pVHL-mediated TCF7L2 degradation, we endeavored to establish which region is vital for the interaction between pVHL and the 26S proteasome. The *in vitro* pulldown assays showed that neither recombinant GST-pVHL(54-99) nor GST-pVHL(100-157) binds to the 19S regulatory subunit RPN10 in purified human 26S proteasome. In contrast, GST-pVHL(54-157) does bind to the 19S regulatory subunit RPN10 (Figure 5C). These results implied that pVHL directly interacts with the 26S proteasome and that pVHL(54-157) alone suffices for this binding. Taken together, these results suggested that the pVHL forms a complex between RPN10 of the 26S proteasome and the TCF/LEFs to mediate TCF/LEF degradation (Figure 5D).

**Figure 5.**
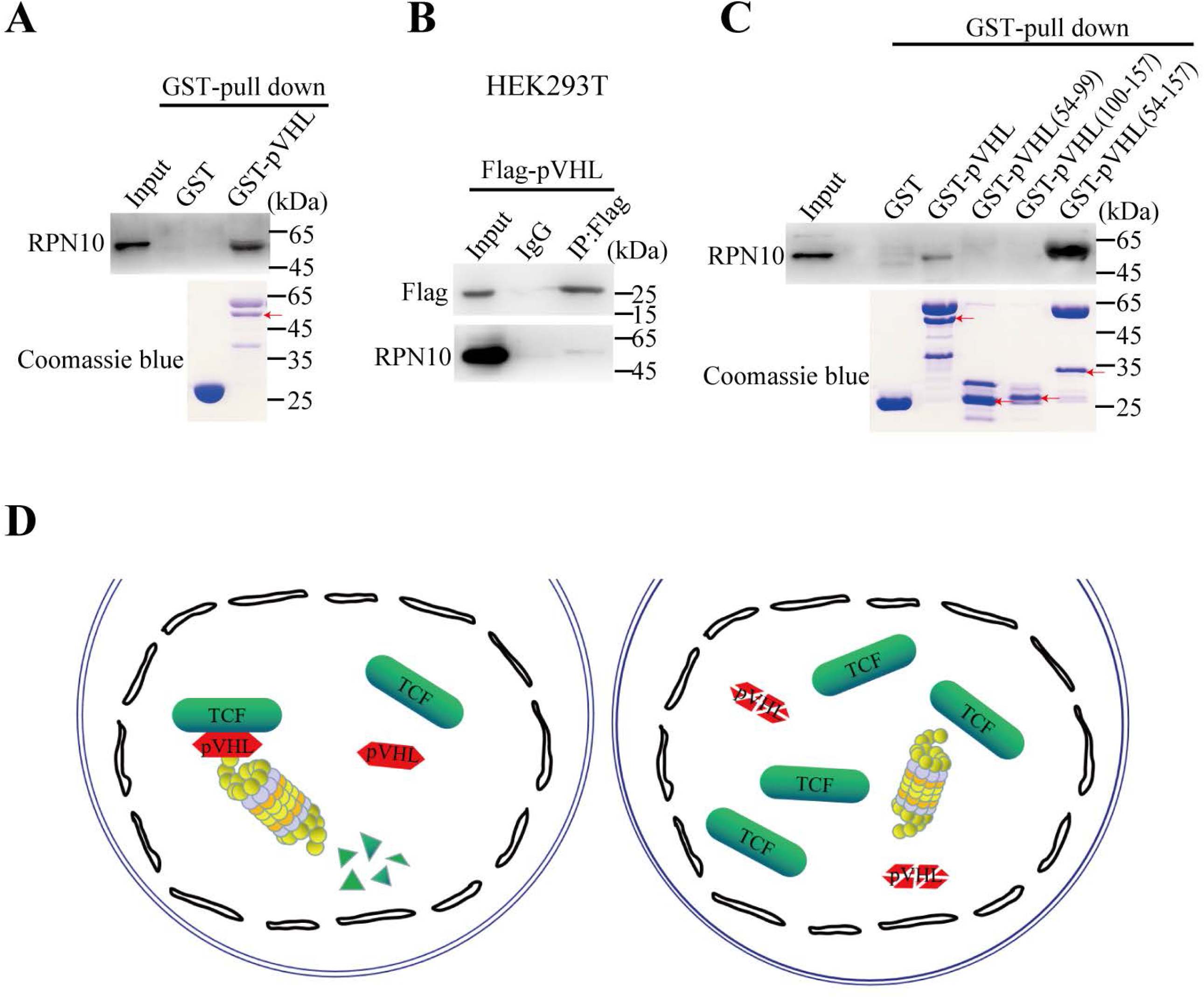
pVHL directly interacts with 26S proteasome. (A) *In vitro* cell-free GST pulldown assay revealed direct interaction between RPN10 and pVHL. Proteasome was incubated with GST or GST-pVHL and pulled down by glutathione agarose. RPN10 is a critical proteasomal subunit protein and was assessed by western blotting. Input proteins were examined by Coomassie blue staining. Red arrow indicates GST-pVHL. (B) Co-IP assay revealed that Flag-pVHL interacted with endogenous RPN10 in HEK293T cells. (C) *In vitro* cell-free GST pulldown assay detected binding of full-length and mutant pVHL to proteasome. Red arrows indicate GST-fusion protein. (D) Schematic illustration of mechanism by which TCF/LEF protein stability is regulated by pVHL.

### pVhl functions upstream of Tcf7l2 to support dHbl and dHbm character

*Vhl*-knockout mice were embryonic lethal and died between E11.5 and 12.5 because of hemorrhagic lesions in the placenta(Gnarra et al., 1997). This limited us to investigate physiological function of Tcf/Lef protein degradation by pVhl. To further uncover the effect of pVHL on TCF/LEF protein degradation *in vivo*, we used a zebrafish *vhl*-null line, which was generated by a CRISPR/Cas9-based gene editing approach(Du et al., 2015). Wnt/β-catenin signaling influences axis formation, including anteroposterior, dorsoventral, and left-right body axis, in vertebrates(Petersen and Reddien, 2009). The *vhl*-null mutant embryos were morphologically indistinguishable from sibling embryos at 48 hpf, while a slightly reduced anterior head end in *vhl*^-/-^ mutant embryos was observed at 96 hpf (Figure 6-figure supplement 1A and D). To further confirm this result, we measured the body length of embryos at 48 hpf and 96 hpf (Figure 6-figure supplement 1B, E). At 48 hpf, the body length of sibling and mutant embryos showed no significant difference (Figure 6-figure supplement 1C). At 96 hpf, the head length in *vhl*-null embryos was shorter than that in sibling embryos, while the trunk length did not exhibit any difference (Figure 6-figure supplement 1F). These results suggested that pVhl is unlikely to play any role in directing dorsoventral or anteroposterior axis formation.

Furthermore, left-right asymmetry and laterality of the heart, visceral organ, and brain were examined. Almost all of the offspring of *vhl* heterozygous mutant showed normal expression of heart marker *cmlc2* at 28 and 48 hpf (Figure 6-figure supplement 2A and B), normal expression of liver and pancreas marker *foxa3* at 48 hpf (Figure 6-figure supplement 2C), and normal expression of liver marker *cp* at 48 hpf (Figure 6-figure supplement 2D). Consistently, the expressions of Nodal ligand *spaw* in the lateral plate mesoderm (LPM), Nodal target genes *lefty1* and *pitx2* in the epithalamus, *pitx2* in the posterior left LPM, and *lefty2* in the heart primordia were not altered at the 23-somite stage (Figure 6-figure supplement 2E-G). These results suggested that depletion of pVhl does not affect expression of left-side Nodal signaling and subsequent organ positioning.

Knockout of pVHL increases TCF/LEF protein levels and enhances Wnt/β-catenin signaling *in vitro*. Additionally, the zebrafish *vhl*-null mutant embryos exhibit increased Tcf7l2 protein levels. In particular, Wnt ligands, Axin1/2, and Tcf7l2 are expressed in the diencephalon and regulate dorsal habenula development(Carl *et al*., 2007; Husken *et al*., 2014; Kuan et al., 2015). Therefore, we next tested whether depletion of pVhl affects Tcf7l2 protein levels in dorsal habenula and habenular neuron development in zebrafish embryos. We monitored the expression of Tcf7l2 on the left and right sides of dHb neurons at 37 hpf, which is immediately after initial Tcf7l2 protein expression in dHb neurons(Husken *et al*., 2014). The *vhl*-null mutant embryos exhibit increased numbers of Tcf7l2^+^ cells on both the left and right sides of dHb neurons (Figure 6A, B), further suggesting that depletion of pVhl increases expression of Tcf7l2 protein. Moreover, we observed that, at 48 hpf, depletion of pVhl strongly reduced the numbers of GFP^+^ cells in *vhl-*null mutant embryos with a *Tg (huc:GFP)* transgenic background (Figure 6C, D). This phenotype did not result from growth retardation since body length, the quantitative indicator of developmental rate, showed no differences between the wild-type and *vhl*-null groups (Figure 6-figure supplement 1A-C). We noticed that this phenotype is reminiscent of embryos after premature activation of Wnt/β-catenin signaling between 24 to 26 hpf, which also delays habenular neuron differentiation with reduced the numbers of GFP^+^ cells in embryos at this stage(Guglielmi *et al*., 2020).

**Figure 6.**
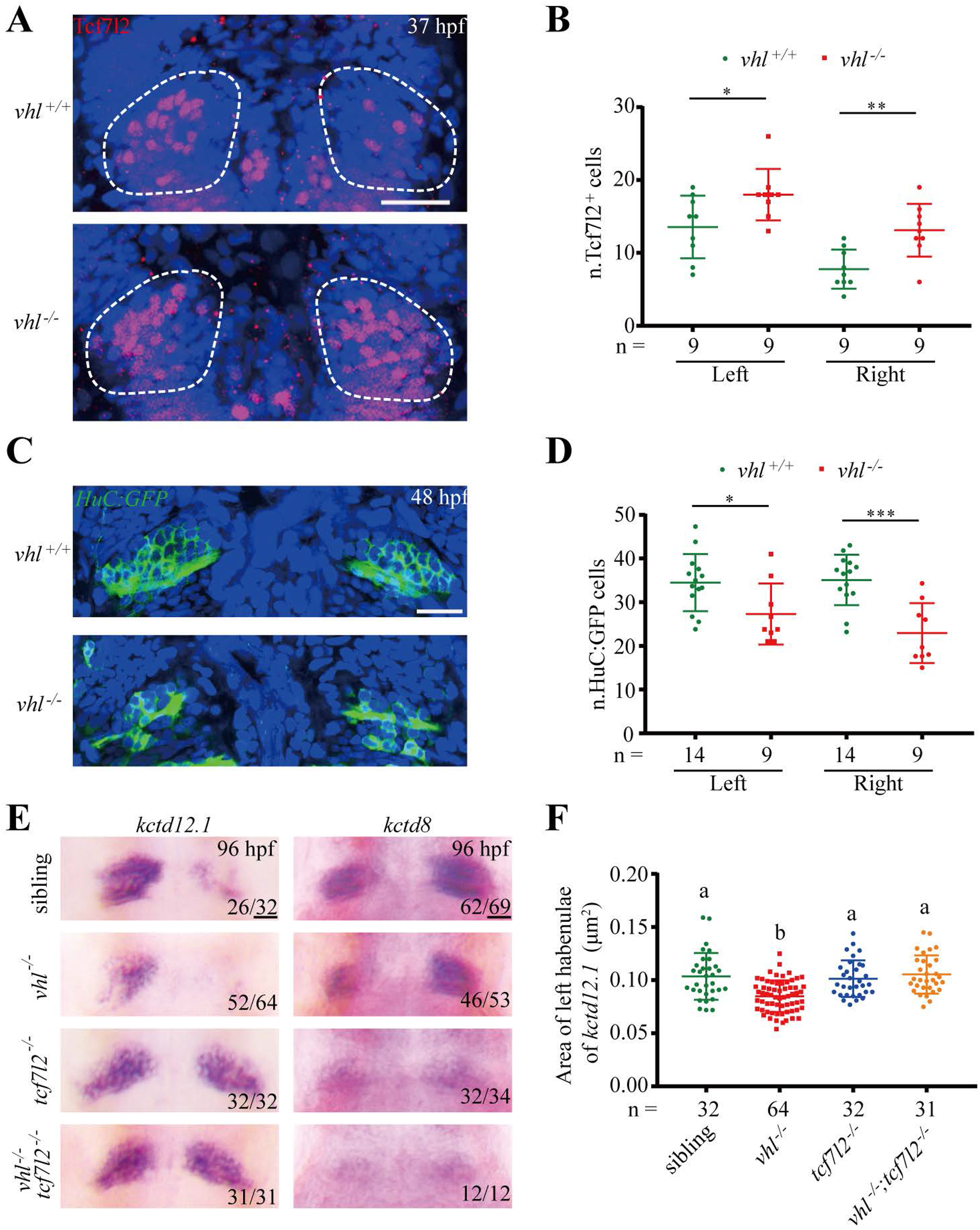
Genetic deletion of *vhl* reduces development of habenular neurons and acts upstream of *tcf7l2*-null mutation. (A) Immunostaining of Tcf7l2-expressing cells in dHb neurons in wild-type and *vhl*-null embryos at 37 hpf. Dorsal view with anterior side upward. Nuclei are counterstained with DAPI (blue), and the habenular region is encircled. Maximum intensity projection of Z-stack images, which were acquired every 2 μm. Scale bar = 25μm. (B)Numbers of Tcf7l2-expressing cells in dHb neurons in (A). Values are mean ± S.D. Unpaired *t*-test. \**p* < 0.05; \*\**p* < 0.01. (C) Habenular neurons of wild-type and *vhl*-null embryos in *Tg (huc:GFP)* transgenic background at 48 hpf. Dorsal view with anterior side upward. Nuclei are counterstained with DAPI (blue). Scale bar = 25 μm. (D) Numbers of left and right lateral HuC:GFP^*+*^ habenular neurons in (C). Values are mean ± S.D. Unpaired *t*-test. \**p* < 0.05; \*\*\**p* < 0.001. (E) Expression of *kctd12*.*1* and *kctd8* in dorsal habenula in embryos with indicated genotypes at 96 hpf. *kctd12*.*1*, which is reduced in *vhl*-null embryos, is enhanced in *tcf7l2*-null mutants, and *kctd8*, which is less strongly reduced in *vhl*-null embryos, is absent in *tcf7l2*-null mutants; *vhl/tcf7l2* double mutants show the same character as that of *tcf7l2*-null mutants. Dorsal view with anterior side upward. Scale bar = 20 μm. (F) Quantification of left lateral *kctd12*.*1*^*+*^ habenular cells in (E). The total embryo numbers are given along the X-axis. Values are mean ± S.D. One-way ANOVA analysis with Tukey’s post-hoc test. Different letters indicate significant differences (*p* < 0.001).

The increased expression of Tcf7l2 protein in dHb neurons may function in premature activation of Wnt/β-catenin signaling and then lead to delayed habenular neuron differentiation. To test this, we examined the epistatic relationship between null mutations in *vhl* and *tcf7l2* by inspecting the expression of dHbl marker *kctd12*.*1* and dHbm marker *kctd8* in progeny embryos of *vhl/tcf7l2* double heterozygous mutants at 96 hpf. Compared with that in wild-type sibling embryos, expression of *kctd12*.*1* was strongly reduced, while that of *kctd8* was less strongly reduced in *vhl*-null embryos (Figure 6E), which was consistent with expression in 96 hpf embryos treated with LiCl or BIO between 24 and 26 hpf(Guglielmi *et al*., 2020). Consistent with the findings of a previous study(Husken *et al*., 2014), *tcf7l2*-null mutant embryos showed enhanced expression of *kctd12*.*1* and reduced expression of *kctd8* (Figure 6E). Considering that pVhl promotes Tcf7l2 proteasomal degradation, pVhl should act upstream of Tcfl2. If the phenotype of *vhl*^*-/-*^ mutants depends on enhanced expression of Tcf7l2 protein, the strongly reduced *kctd12*.*1* expression and less strongly reduced *kctd8* expression should not be observed in *vhl/tcf7l2* double mutants, instead of the phenotype of *tcf7l2* single mutant. In other words, the expression of *kctd12*.*1* and *kctd8* in a *tcf7l2*^*-/-*^ background should be enhanced and reduced respectively, irrespective of the presence or absence of pVhl function. As expected, the expression of *kctd12*.*1* and *kctd8* in *vhl/tcf7l2* double mutants was consistent with that in *tcf7l2* single mutant embryos (Figure 6E). We quantified the expression area of left lateral *kctd12*.*1* in Figure 6E. Significant reduction was only observed in *vhl* single mutant embryos (Figure 6F). Collectively, these results suggested that pVhl likely acts upstream of Tcf7l2 and plays a specific role in dorsal habenula development.

## Discussion

The TCF/LEF transcription factor family includes four members that display distinct and sometimes redundant functions. In this study, we discovered that pVHL directly interacts with all four TCF/LEF family members and promotes their degradation via a ubiquitin-independent proteasomal degradation mechanism. In this way, pVHL regulates TCF/LEF protein abundance and may therefore be crucial in the Wnt/β-catenin signaling cascade in the nucleus, which ensures elaborate and temporal control of the output of this signaling. Using *vhl-*and *tcf7l2-*knockout zebrafish, we found that pVhl functions in regulating the development of dorsal habenular neurons in zebrafish embryos and likely acts by modulating the protein stability of Tcf7l2.

It has been reported that Lef1, Tcf7, and Tcf7l2 act as β-catenin-dependent transcriptional activators, whereas Tcf7l1 functions as a transcriptional repressor(Cole et al., 2008; Kim et al., 2000; Merrill et al., 2004; Yi et al., 2011). In addition, Wnt activation promotes β-catenin binding and inactivates Tcf7l1 but not Lef1, Tcf7, or Tcf7l2 by reducing the chromatin occupancy of Lef1 and secondarily stimulates Lef1 protein degradation by proteasome in embryonic stem cells, but the underlying mechanism is not clear(Shy *et al*., 2013). Regulation of TCF/LEF protein stability is an important but poorly understood issue. Ishitani et al. (2005) reported that MG132 treatment stabilizes LEF1 and increases its ubiquitination level, while deletion of the LEF1 C-terminus reduces ubiquitination to a lesser extent. LEF1 is degraded by the ubiquitin-proteasome pathway, and its ubiquitination sites may be located in its C-terminal region(Ishitani *et al*., 2005). Yamada et al. (2006) reported that the E3 ubiquitin-ligase NARF ubiquitylates TCF7L2 and LEF1 and promotes their degradation via the proteasome pathway. These findings suggest that some TCF/LEF protein stability is regulated and also degraded via the ubiquitin-proteasome pathway. Recently, we found that VBP1, a pVHL binding protein, promotes TCF/LEFs protein proteasomal degradation when pVHL is present(Zhang *et al*., 2020). In this study, we further demonstrated that pVHL promotes the degradation of all four TCF/LEF members. Intriguingly, this function of pVHL is not dependent on its E3 ubiquitin ligase activity but on an unexpected ubiquitin-independent proteasome pathway. This discovery suggested that TCF/LEF can undergo the ubiquitin-independent proteasome pathway.

We made several findings in this study: First, TCF/LEF transcription factors are pVHL substrates. In contrast, pVHL binds to its canonical substrates HIF-α, Akt, and ZHX2, which undergo prolyl hydroxylation and are recognized by the hydroxyl-proline binding pocket in pVHL. TCF7L2 is degraded by pVHL as well as by pVHL with mutated hydroxyl-proline binding pocket sites. Prolyl hydroxylase inhibitor had little effect on the promotion of TCF7L2 degradation by pVHL. Therefore, TCF/LEFs are degraded by pVHL without prolyl hydroxylation. Second, pVHL promotes TCF/LEF degradation via its aa 54-157 region in a manner independent of E3 ubiquitin ligase activity. The aa 54-157 region is necessary and sufficient for TCF/LEF degradation by pVHL. Within this region, the aa 100-157 region is critical for pVHL binding to TCF7L2. pVHL promotes TCF/LEF degradation in both HEK293T cells and HCT116 cells. In addition, we observed that depletion of pVHL in HEK293T cells amplified BIO treatment-induced Wnt reporter activity. Therefore, pVHL promotes the proteasomal degradation of TCF/LEFs in the Wnt-off and Wnt-on states. Third, pVHL promotes TCF/LEF proteasomal degradation via a heretofore unknown ubiquitin-independent pathway, and pVHL (54-157) binds directly to the 19S regulatory subunit RPN10 of the 26S proteasome.

Another interesting observation made in this study is that pVhl regulates the development of dHb neurons in zebrafish embryos likely by modulating the protein stability of Tcf7l2. Knockout of pVhl leads to a reduction in dHb neurons, which was evidenced not only by a specific reduction in the number of HuC:GFP^+^ cells in differentiating habenular neurons but also by a reduction in expression area of indicative markers of dHbl and dHbm at later stages; however, organ positioning and left-side Nodal signaling in *vhl*-null mutants were not altered. This is different from the phenotype of *axin1* mutant embryos, which exhibit bilateral expression of the Nodal pathway genes in the epithalamus and unaffected left-side Nodal signaling and organ positioning within the LPM(Carl *et al*., 2007). In contrast, the above phenotype in *vhl*-null mutants produces phenocopied embryos after treatment with LiCl or BIO between 24 and 26 hpf, leading to premature activation of Wnt signaling(Guglielmi *et al*., 2020). Overactivation of Wnt/β-catenin signaling in the gastrulation or mid-somite stage disrupts the laterality of Nodal pathway expression in both the LPM and brain(Carl *et al*., 2007; Lin and Xu, 2009). However, these effects were not observed in *vhl*-null mutants. Thus, it seems unlikely that the phenotype in *vhl*-null mutants results from the possibly increased activity of Wnt/β-catenin signaling in the gastrulation or mid-somite stage by depletion of pVhl. Alternatively, this phenotype may arise from the initial increased Tcf7l2 expression in dHb neurons, which causes temporally and locally increased Wnt activity. Future studies will be needed to determine the comprehensive developmental basis underlying the phenotype after depletion of zebrafish pVhl. Indeed, the *vhl*-null mutation is upstream of functional Tcf7l2, as shown by an epistatic analysis between *vhl*-null and *tcf7l2*-null mutations. As mentioned earlier, TCF/LEF family in vertebrates contains four members. Disruption of any one results in a distinct phenotype(Cadigan and Waterman, 2012). Future efforts are needed to investigate the potential physiological effect of interaction between pVHL and distinct member of TCF/LEF family.

In summary, our study identified that TCF/LEF is degraded by pVHL via a ubiquitin-independent proteasomal degradation pathway. Importantly, we further showed that pVhl is involved in the development of dHb neurons in zebrafish embryos and likely functions by promoting the degradation of Tcf7l2. Therefore, regulation of the stability of TCF/LEF protein is as important as that of β-catenin protein in Wnt signaling pathway, as both of them control the Wnt signaling output. pVHL regulates the degradation of DVL, β-catenin, and TCF/LEFs, suggesting that pVHL likely plays an essential role in controlling the Wnt signaling output by regulating the protein levels of the key components of Wnt signaling at different signal levels. An earlier study suggested that the promoter of *VHL* responds to β-catenin/TCF7L2 and that pVHL has interplay with the Wnt/β-catenin pathway during colorectal tumorigenesis(Giles et al., 2006). Our findings identify a mechanistic connection between these two important signaling pathways and may facilitate deeper understanding of the interplay between the pVHL and Wnt/β-catenin signaling pathways in embryogenesis, organogenesis, and even diseases. Moreover, pVHL is a well-known tumor suppressor, as up to 92% of clear cell renal carcinomas have an inactivated *VHL* gene. Upregulated Wnt/β-catenin signaling might also contribute to renal carcinogenesis associated with *VHL* mutations(Saini et al., 2011). These findings made herein link pVHL to oncogenic Wnt/β-catenin signaling at the TCF/LEF level. The efforts to determine that pVHL mediates substrate degradation by direct interaction and complexation with the 26S proteasome should have broad utility for better understanding the physiological role and molecular function of pVHL in the future.

## Materials and Methods

### Chemicals, reagents, and antibodies

Dulbecco’s modified Eagle’s medium (DMEM) was purchased from Hyclone (Logan, UT, USA). Fetal bovine serum (FBS) was purchased from PAN (Aidenbach, Germany). Protein A/G Plus-agarose was purchased from Santa Cruz Biotechnology (Dallas, TX, USA). The antibodies used included the following: rabbit anti-TCF7L2 (1:1,000 for western blotting and 2 μg for co-IP assays; #2569; Cell Signaling Technology, Danvers, MA, USA), rabbit anti-TCF7 (1:1,000 for western blotting; #2203; Cell Signaling Technology,), rabbit anti-TCF7L1 (1:1,000 for western blotting; #2883; Cell Signaling Technology, Danvers, MA, USA), mouse anti-TCF7L1/TCF7L2 (1:500 for whole mount immunohistochemical staining; #ab12065; Abcam, Cambridge, UK), rabbit anti-pVHL (1:1,000 for western blotting, #68547; Cell Signaling Technology), mouse anti-pVHL (1:1,000 for western blotting, sc-135657; Santa Cruz Biotechnology), rabbit anti-proteasome 19S S5A (1:1,000 for western blotting; ab137109; Abcam, Cambridge, UK), mouse anti-HIF-1α (1:1,000 for western blotting; 610958, BDBioscience, Franklin Lakes, NJ, USA), rabbit anti-HIF-2α (1:1,000 for western blotting; NB 100-122; Novus Biologicals, Centennial, CO, USA), rabbit anti-HIF-1β (1:1,000 for western blotting; A19532, ABclonal, Wuhan, China), rabbit anti-GAPDH (1:1,000 for western blotting; D110016; BBI, Crumlin, UK), rabbit anti-Histone H3.1 (1:1,000 for western blotting; #P30266; Abmart, Shanghai, China), mouse anti-Myc (1:1,000 for western blotting and 2 μg for co-IP assays; sc-40; Santa Cruz Biotechnology), and mouse anti-Flag (1:1,000 for western blotting and 2 μg for co-IP assays; F1804; Sigma, St. Louis, MO, USA). Primers and sequence information are provided in Supplementary Table S1.

### Molecular cloning and plasmid construction

The plasmids pCS2-6×Myc-Tcf7, pCS2-6×Myc-Tcf7l1, pCS2-6×Myc-Tcf7l2, and pCS2-6×Myc-Lef1 were gifts from Dr. Wei Wu (School of Life Sciences, Tsinghua University, China). pCS2-Flag-pVHL, pGEX-2T-pVHL, pGEX-2T-pVHL(54-99), pGEX-2T-pVHL(100-157), pGEX-2T-pVHL(54-157), pCS2-VP16-Tcf7l1ΔN, and pCDNA3-HA-Ub, pBoBi-puro-GFP, pBoBi-puro-Flag-pVHL, pCS2-pVHL-P2A-GFP, and pCS2-pVHL(1-157)-P2A-GFP were generated by PCR subcloning. The VHL mutants pVHLΔ(1-53), pVHLΔ(54-99), pVHLΔ(100-157), pVHLΔ(158-213), pVHL(1-157), pVHL(54-99), pVHL(100-157), and pVHL(54-157) were amplified by PCR and subcloned into pCS2-Flag or pCS2-eGFP. The pCS2-Flag-pVHL L158P and pCS2-Flag-pVHL R167W mutants were generated by site-directed mutagenesis.

### Zebrafish strains

Zebrafish (*Danio rerio*) Tübingen wild-type, the transgenic line *Tg*(*huc*:*GFP*), and the *vhl*-null and *tcf7l2*-null strains were maintained on a 14h light/10h dark cycle at 28.5 °C and fed twice daily. The zebrafish *vhl* mutant strain was gifted by Dr. Wuhan Xiao(Du *et al*., 2015). The zebrafish *tcf7l2* mutant allele (*tcf7l2*^+/ihb316^, ZFIN ID: ZDB-ALT-181129-11) was generated by the CRISPR/Cas9 method and obtained from the China Zebrafish Resource Center, National Aquatic Biological Resource Center (CZRC/NABRC), Wuhan, China. The animals were raised and maintained according to standard procedures described in Zebrafish Information Network (ZFIN; https://zfin.org/). Embryos obtained by natural crosses were maintained in embryo rearing solution in an incubator at 28.5°C. The embryos were staged according to standard methods(Kimmel et al., 1995). All experimental protocols were approved by and conducted in accordance with the Ethical Committee of Experimental Animal Care, Ocean University of China.

### Cell lines and transfections

HEK293T, HeLa, HCT116, and 786-O cell lines were purchased from the American Type Culture Collection (ATCC; Manassas, VA, USA). HEK293T, HeLa, and HCT116 cells were cultured in DMEM supplemented with 10% FBS and 1% penicillin/streptomycin at 37 °C in 5% CO_2_. The 786-O cells were cultured in RPMI medium supplemented with 10% FBS and 1% penicillin/streptomycin at 37 °C in 5% CO_2_. Hypoxia-treated cells were performed in a hypoxic chamber containing 1% O_2_ at 37 °C for 24 h as reported previously(Zhang et al., 2014). For starvation treatment, HEK293T cells were washed twice with DMEM and cultured under serum starvation in a time series. In some experiments, MG132 (10 μM) was used to treat cells for 8 h to inhibit proteasome activity, or NH_4_Cl (25 mM) was used to disrupt lysosome function, or 3-MA (5 mM) was used to inhibit autophagy. DMOG (200 μM) was added to cultured HEK293T cells for 12 h to antagonize prolyl hydroxylase. Plasmid transfection was performed using polyethylenimine (#23966-2; Polysciences Inc., Warrington, PA, USA) following the manufacturer’s instructions. To generate Flag-pVHL stable cells, packaging of lentiviral *VHL* cDNA expressing viruses and subsequent infection of 786-O cell line were performed. Following viral infection, cells were maintained in the presence of puromycin (1 μg/mL). Knock-in efficiency of Flag-pVHL was examined by immunoblotting assay.

### GST fusion protein purification and *in vitro* GST pulldown assays

GST-pVHL, GST-pVHL(54-99), GST-pVHL(100-157) and GST-pVHL(54-157) were expressed in *E. coli* BL21. The fermentation broth was started with 1% inoculant incubated at 37°C with agitation at 150 rpm until OD_600_ = 0.6 ∼ 0.8. Fusion proteins were induced with 1 mM IPTG at 37 °C for 5 h. Cells were harvested by centrifugation, washed with ice-cold phosphate-buffered saline (PBS), and lysed by sonication in lysis buffer (50 mM Tris at pH 7.5, 150 mM NaCl, 1 mM EDTA, 10% glycerol, and 1% Triton X-100) on ice. The GST and GST fusion protein extracts were mixed with glutathione Sepharose 4B beads (71024800-GE; GE Healthcare, Chicago, IL, USA) overnight at 4 °C and the mixtures were then washed three times with ice-cold PBS. To test the direct interaction between pVHL and Tcfs, the mixtures were centrifuged, and the precipitates were diluted with equal volumes of the indicated HEK293T cell lysates and mixed by rotation overnight at 4 °C. After extensive washing with lysis buffer, the precipitated proteins were eluted by SDS/PAGE. Bound proteins were detected either with the indicated antibodies or by Coomassie blue staining. To test the direct interaction between pVHL and proteasome, the bait GST or GST fusion protein was incubated with purified human 26S proteasome (E365; Boston Biochem, Cambridge, MA, USA) at 4 °C for 2 h in binding buffer (1× PBS, 2 mM EDTA, 1 mM phenylmethylsulfonyl fluoride, and 0.5% Triton X-100). After washed three times, and the bound proteins were analyzed by western blot.

### Luciferase assays

The cells were transfected with TOPFlash reporter. Transfection efficiency was normalized by co-transfection with a *Renilla* reporter. Cells were lysed with 1× passive lysis buffer (Promega, Madison, WI, USA). TOPFlash/*Renilla* luciferase assays were performed using the dual-luciferase reporter assay kit (Promega) according to the manufacturer’s instructions.

### Cycloheximide chase assay

HEK293T cells were co-transfected with Myc-Tcf7l2 and an empty vector, Flag-pVHL or Flag-pVHL (1-157). After 16 h, the cells were treated with CHX (100 μg/mL) and harvested at the indicated time points. Lysates were prepared and subjected to immunoblot analysis. Protein stability was determined from the percentage of GAPDH-normalized Tcf7l2 remaining at an indicated point relative to the initial time point.

### Ubiquitination assay

Ubiquitination assays were performed using hot lysis-extracted protein lysates according to a described protocol(Du et al., 2016). Briefly, HEK293T cells were treated with 10 μM MG132 for 8 h before being harvested, and the cells were hot-lysed by boiling in 100 μL denaturing buffer (2% SDS, 10 mM Tris-HCl [pH 8.0], 150mM NaCl, and 1× protease inhibitor mixture with 10 mM freshly prepared N-ethylmaleimide) for 10 min. The lysates were diluted ten-fold with dilution buffer (1% Triton X-100, 10 mM Tris-HCl [pH 8.0], 150 mM NaCl, 2 mM EDTA, and 1× protease inhibitor mixture with 10 mM freshly prepared N-ethylmaleimide). Tcf7l2 and HIF-1α were immunoprecipitated from whole-cell lysate by incubating it with the appropriate antibodies and Protein A/G resin. Ubiquitinated Tcf7l2 and HIF-1α were detected by immunoblot using the indicated antibodies.

### Western blot and immunoprecipitation

Cells or zebrafish embryos were lysed in RIPA buffer (150 mM NaCl, 1% Triton X-100, 1% sodium deoxycholate, 0.1%SDS, and 50 mM Tris at pH 7.5) supplemented with protease and phosphatase inhibitors on ice for 15 min and centrifuged at 12,000 rpm at 4 °C for 10 min. Protein samples were separated by SDS/PAGE and transferred to PVDF membranes. For the Co-IP experiments, the cells were lysed in IP lysis buffer containing 50 mM Tris (pH 7.5), 150 mM NaCl, 1 mM EDTA, 10% glycerol, 1% Triton X-100, and protease and phosphatase inhibitors. Cleared cell lysates were incubated with the appropriate antibodies (1-2 μg) overnight at 4 °C followed by 4 h incubation at 4 °C with Protein A/G agarose beads. All immune complexes bound to the Protein A/G beads were washed five times with IP lysis buffer and detached from the agarose with SDS loading buffer for immunoblot analysis. A minimum of three independent experiments were performed.

### Immunofluorescence staining

Cells were grown on coverslips. After transfection, the cells were fixed with 4% paraformaldehyde, permeabilized for 5 min at room temperature with PBS containing 0.25% Triton X-100, and blocked with 3% bovine serum albumin in PBS. The cells were incubated with anti-Myc antibody overnight at 4 °C. Goat anti-mouse Cy3 was used as the secondary antibody. The cells were mounted in mounting medium after incubation with PBS containing DAPI. Images were acquired and viewed with use of a Leica SP8 confocal microscope (Leica Microsystems, Wetzlar, Germany).

### Generation of knockout cell lines via CRISPR/Cas9

The gRNAs were designed in CRISPR V2 (http://zifit.partners.org). The sequence (5′-CGCGTCGTGCTGCCCGTAT-3′) in the first exon was chosen as the CRISPR targeting site for *VHL* in the human cell line. The *ARNT* target sequence in the sixth exon was 5′-TGAAATTGAACGGCGGCGA-3′. For CRISPR knockout, cells were transfected with plasmids expressing indicated gRNA and Cas9 and selected by antibiotics.

### Quantitative real-time RT-PCR

Total RNA was isolated from cultured cells using TRIzol reagent (Invitrogen, Carlsbad, CA, USA). The cDNAs were reverse-transcribed into first-strand cDNA using Oligo(dT)_18_ and M-MLV according to the manufacturer’s instructions. Quantitative real-time RT-PCR (qRT-PCR) was performed using an iCycler iQ Multicolor Real-time PCR Detection System (Bio-Rad Laboratories, Hercules, CA, USA). Primer sequences used for the PCR experiments were as reported previously(Zhang *et al*., 2020). Samples from three independent experiments were collected, and each sample was measured in duplicate. The mRNA levels of the genes of interest were calculated using the 2^-ΔΔCt^ method and normalized to *β-actin*.

### Microinjection

The capped mRNAs were generated *in vitro* with the mMESSAGE mMACHINE Kit (Ambion, Austin, TX, USA). Diluted mRNA was injected into 1-2 cell stage zebrafish embryos. The protein expression of GFP in zebrafish embryos was detected at 6 hpf.

### Whole-mount *in situ* hybridization and immunohistochemical staining

Whole-mount *in situ* hybridization using a digoxigenin-labeled RNA riboprobe was performed as reported previously(Feng et al., 2012). Stained embryos or live embryos were mounted in glycerol or 5% methylcellulose, respectively. Images were taken using a Leica M205 FCA microscope. Image area analysis of the left of the *kctd12*.*1* measurement was performed with Image J software.

Antibody staining was performed according to standard procedures(Turner et al., 2014). Embryos were fixed with formaldehyde at the indicated time points and washed with PBST with 0.8% Triton X-100 in PBS added. Embryos at 36-48 hpf were digested with proteinase K for 30 min and then fixed with formaldehyde for 20 min. After being blocked for at least 1 h, embryos were incubated in the primary antibody in 4 °C overnight. After being washed sufficiently with PBST, embryos were incubated in the second antibody in 4 °C overnight. After being washed, the nuclei stained with DAPI.

### Confocal microscopy and image analysis

Confocal imaging was performed on a Leica TCS SP8 STED microscope. The *Tg*(*huc:GFP*) embryos or whole mount immunohistochemical stained embryos were mounted in 1.2% low-melt agarose in glass-bottom dishes. Images were acquired with a 40× water objective. To count dHb neurons, we used the transgenic lines *Tg*(*huc:GFP*) in combination with nuclear DAPI staining. Left and right HuC:GFP+ neurons were counted using Image J from confocal stacks acquired every 2 μm.

### Statistical analysis

Graphs were plotted with GraphPad Prism 7 Software (GraphPad Software, La Jolla, CA, USA). Statistical analyses were performed using a two-tailed, unpaired Student’s *t*-test for comparisons between two groups or one-way ANOVA analysis of variance followed by Tukey’s Bonferroni’s and Dunnett’s post-hoc test was used for comparisons among multiple groups or Two-way ANOVA analysis of variance followed by Bonferroni’s post-hoc test was used for two independent variables affect a dependent variable. *p*<0.05 or smaller *p* value was considered statistically significant. Unless otherwise indicated, all experiments were performed in triplicate, and the data were reported as means ± S.D. for three experiments.

## Acknowledgements

We are grateful to Dr. Wuhan Xiao from Institute of Hydrobiology, Chinese Academy of Sciences for providing the *vhl* mutant zebrafish. We are grateful to Dr. Matthias Carl from Department of Cellular, Computational and Integrative Biology (CIBIO), University of Trento, for providing advice on distinguishing the habenular regions.

## Financial disclosure

This work was supported by the National Key R & D Program of China (2018YFA0801000 to JZ), the National Natural Science Foundation of China-Shandong Joint Fund (U1606403 to JZ), the Fundamental Research Funds for the Central Universities (201822023 to JZ, 201762022 to XR), the National Natural Science Foundation of China (31601863 to XR, 32170834 to JZ, 31872189 to JZ, and 30972238 to JZ), and the Natural Science Foundation of Shandong Province (ZR2017MC001 to JZ). The funders had no role in the study design, data collection and analysis, decision to publish, or preparation of the manuscript.

## Figure Legends

**Figure 1-figure supplement 1.**
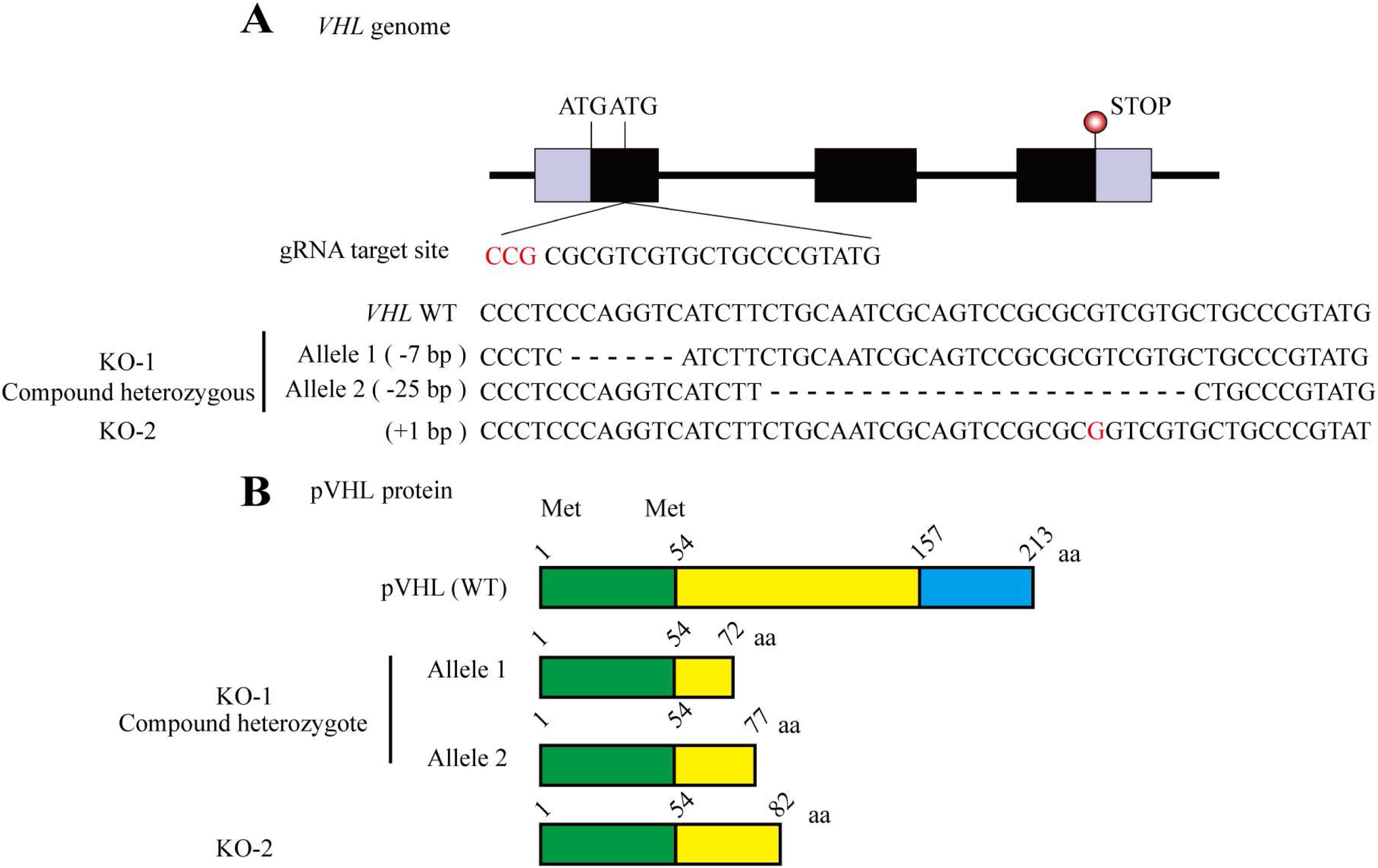
Generation of *VHL*-null cell lines. (A) Schematic illustrations of genomic structures and target positions of CRISPR/Cas9-mediated *VHL* mutation. ATG denotes translation start codon; the black box denotes exon; purple box denotes UTR; black lines denote introns. (B) Schematic illustrations of pVHL truncated protein structures. Two Met denote different translation start codons in pVHL. Numbers denote amino acid positions of critical domain and mutant protein length.

**Figure 2-figure supplement 1.**
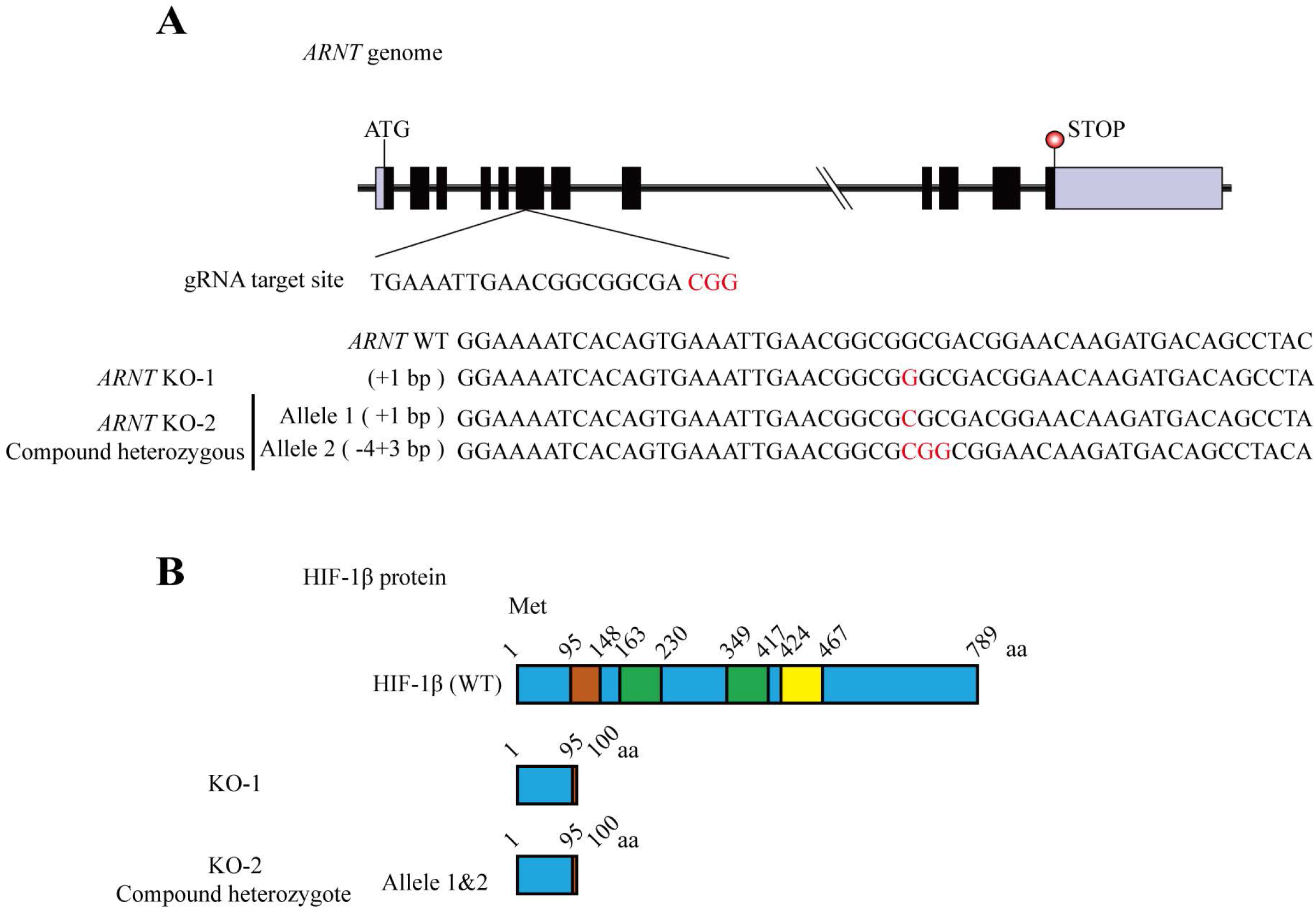
Generation of *HIF1-β (ARNT)*-null cell lines. (A) Schematic illustrations of genomic structures and target positions of CRISPR/Cas9-mediated *ARNT* mutation. ATG denotes translation start codon; the black box denotes exon; purple box denotes UTR; black lines denote introns. (B) Schematic illustrations of HIF1-β truncated protein structures. Numbers denote amino acid positions of critical domain and mutant protein length.

**Figure 2-figure supplement 2.**
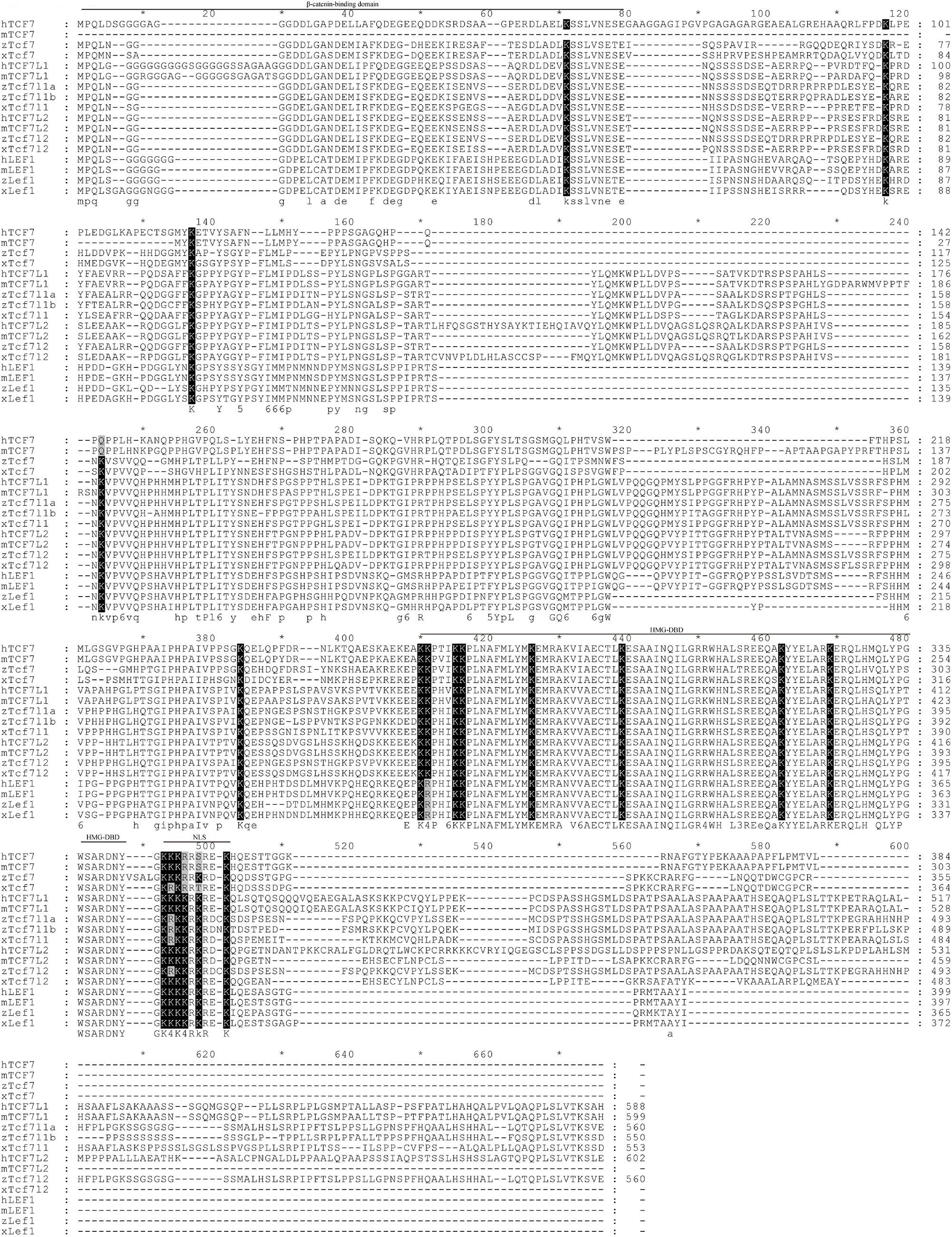
TCF/LEF amino acid sequence alignment. Amino acid sequence alignment of human, mouse, *Xenopus*, and zebrafish TCF/LEFs. Conserved lysines are indicated in black. Accession numbers are: human TCF7 NP_003193.2, mouse TCF7 NP_001300910.1, zebrafish Tcf7 NP_001012389.1, *Xenopus* Tcf7 NP_989421.1, human TCF7L1 NP_112573.1, mouse TCF7L1 NP_001073290.1, zebrafish Tcf7l1a NP_571344.1, zebrafish Tcf7l1b NP_571371.2, *Xenopus* Tcf7l1 NP_001005640.1, human TCF7L2 NP_001139746.1, mouse TCF7L2 NP_001136390.1, zebrafish Tcf7l2 NP_571334.1, *Xenopus* Tcf7l2 NP_001231922.1, human LEF1 NP_057353.1, mouse LEF1 NP_034833.2, zebrafish Lef1 NP_571501.1 and *Xenopus* Lef1 NP_001230763.1.

**Figure 4-figure supplement 1.**
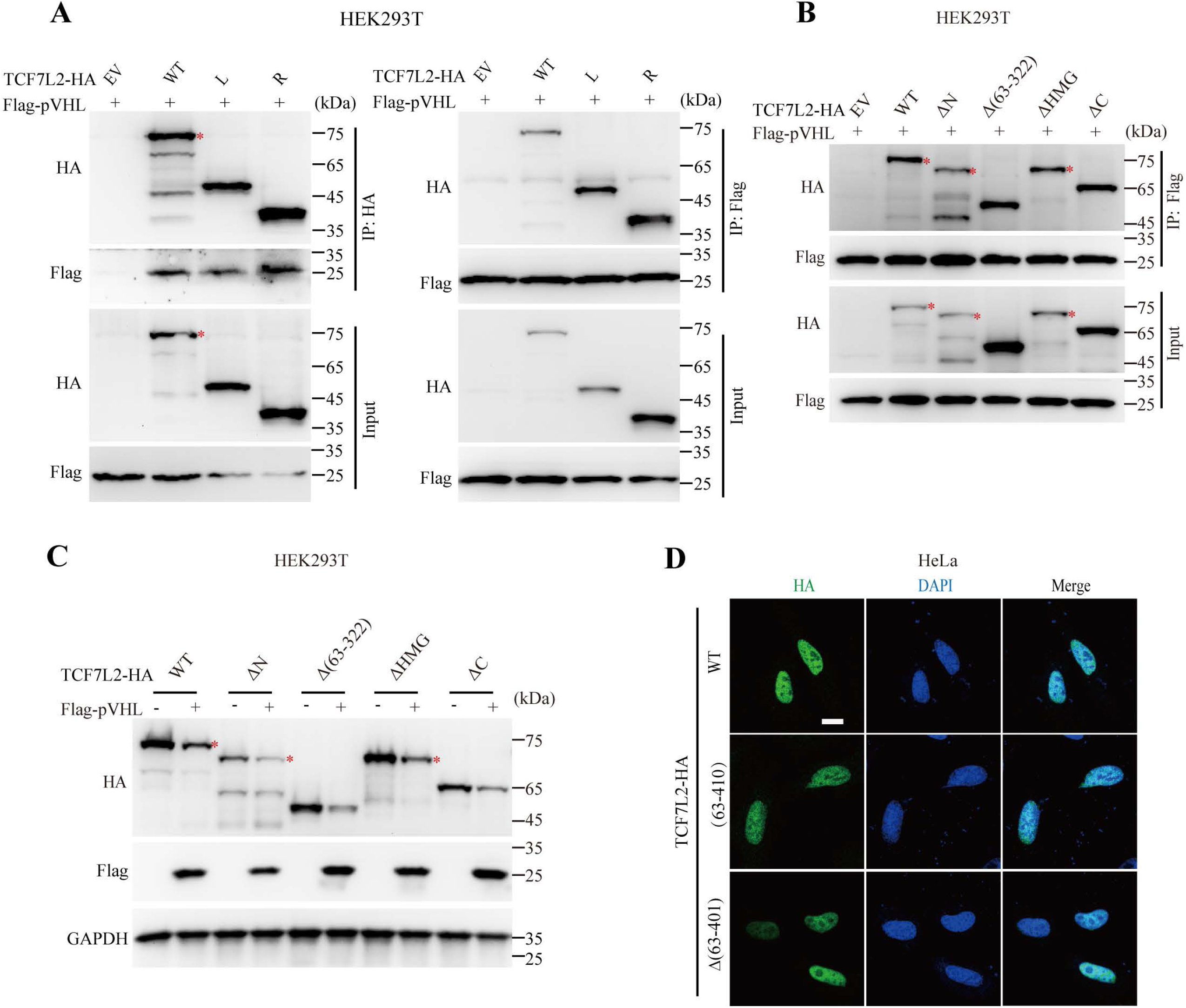
Mapping the binding domains of TCF7L2 to pVHL. (A, B) Mapping TCF7L2 binding domain associated with pVHL in transfected HEK293T cells by Co-IP assay. Red asterisk indicates the specific band. (C) The protein levels of TCF7L2 mutants in HEK293T cells with overexpression of pVHL. (D) The cellular location of HA-tagged TCF7L2 mutants in HeLa cells. Scale bar =10 μm.

**Figure 6-figure supplement 1.**
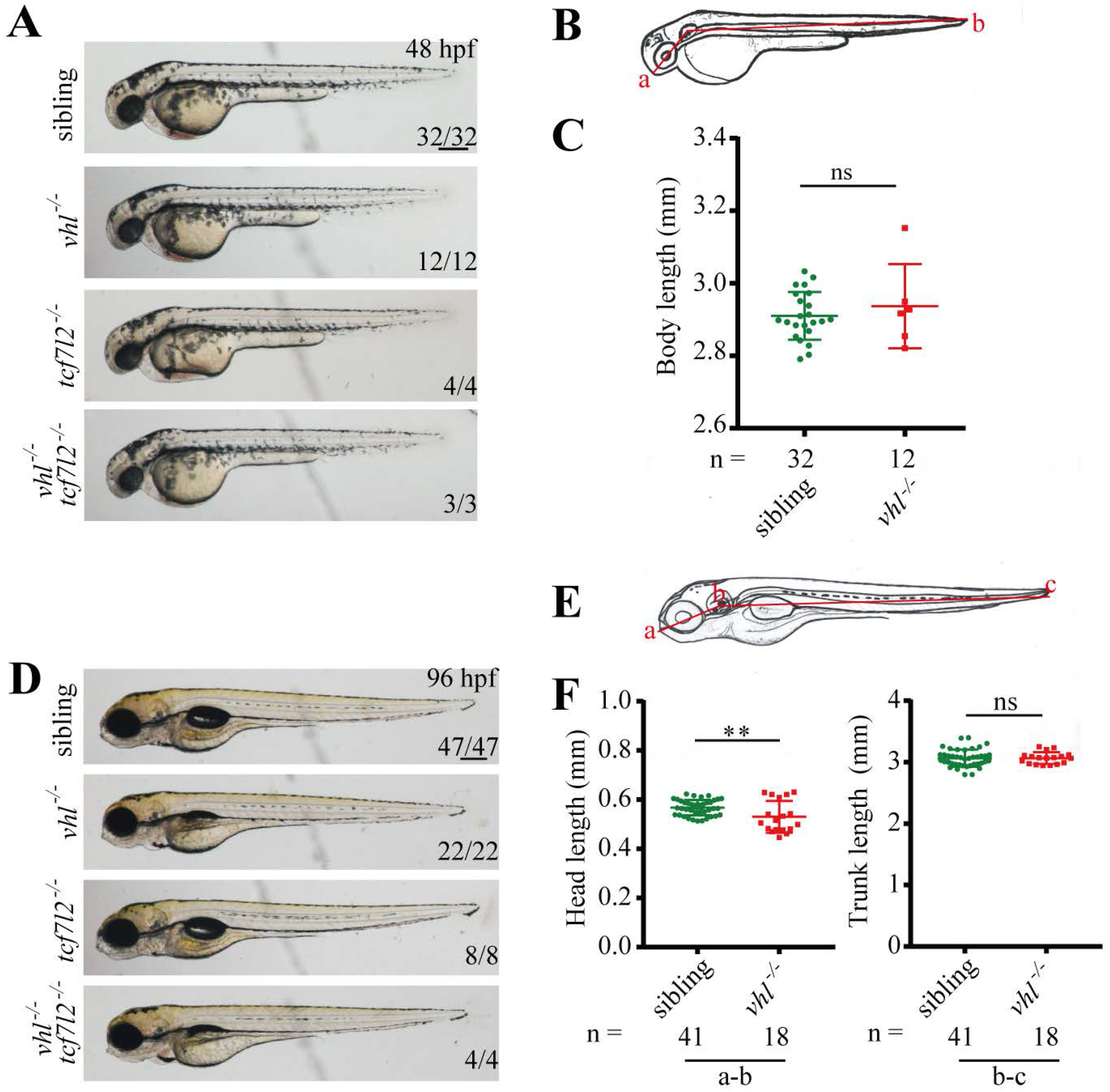
Representative morphologies of *vhl, tcf7l2*, and *vhl/tcf7l2* double mutants. (A, D) Representative images of mutant embryos at 48 hpf (A) and 96 hpf (D) with indicated genotypes. Scale bar = 250 μm. (B, E) Schematic illustration representing body length measurement at 48 hpf (A) and 96 hpf (D). (C) Quantification of the body length of a-b (mouth to end of tail through center of ear vesicle) in sibling and *vhl* mutant embryos at 48 hpf (A). Values are mean ± S.D. Unpaired *t*-test. ns, not significant. (F) Quantification of the head length of a-b (mouth to center of ear vesicle) and trunk length of b-c (center of ear vesicle to end of tail) in sibling and *vhl* mutant embryos at 96 hpf (D). The total embryo numbers are given along the X-axis. Values are mean ± S.D.. Unpaired *t*-test. ns, not significant; ** *p* < 0.01.

**Figure 6-figure supplement 2.**
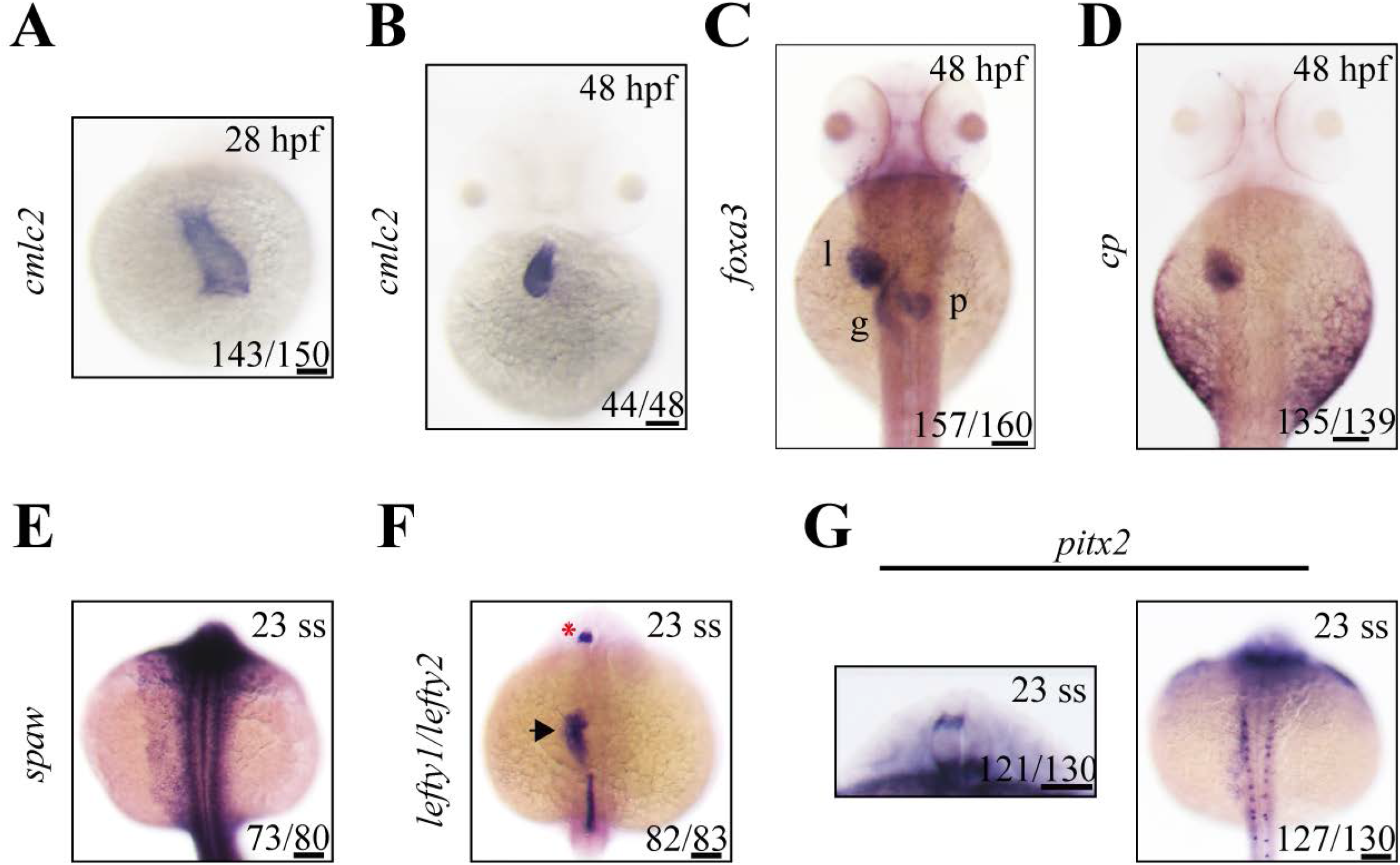
Depletion of pVhl had little effect on left-right asymmetric development. (A-G) Offspring embryos of heterozygous *vhl* mutants are examined for *cmlc2* expression at 28 hpf (A) and 48 hpf (B), *foxa3* expression at 48 hpf (C), *cp* expression at 48 hpf (D), *spaw* expression at the 23-somite stage (E), *lefty1/lefty2* expression at the 23-somite stage (F), *pitx2* expression in head (left) and LPM (right) at the 23-somite stage (G). Embryos are shown in ventral (A, B) or dorsal view (C-G) with anterior side upward. The asterisk indicates the expression of *lefty1* in the diencephalon, and the arrow indicates the expression of *lefty2* in heart field (F). l, liver; p, pancreas; g, gut. Scale bar = 100 μm.

**Table S1.**
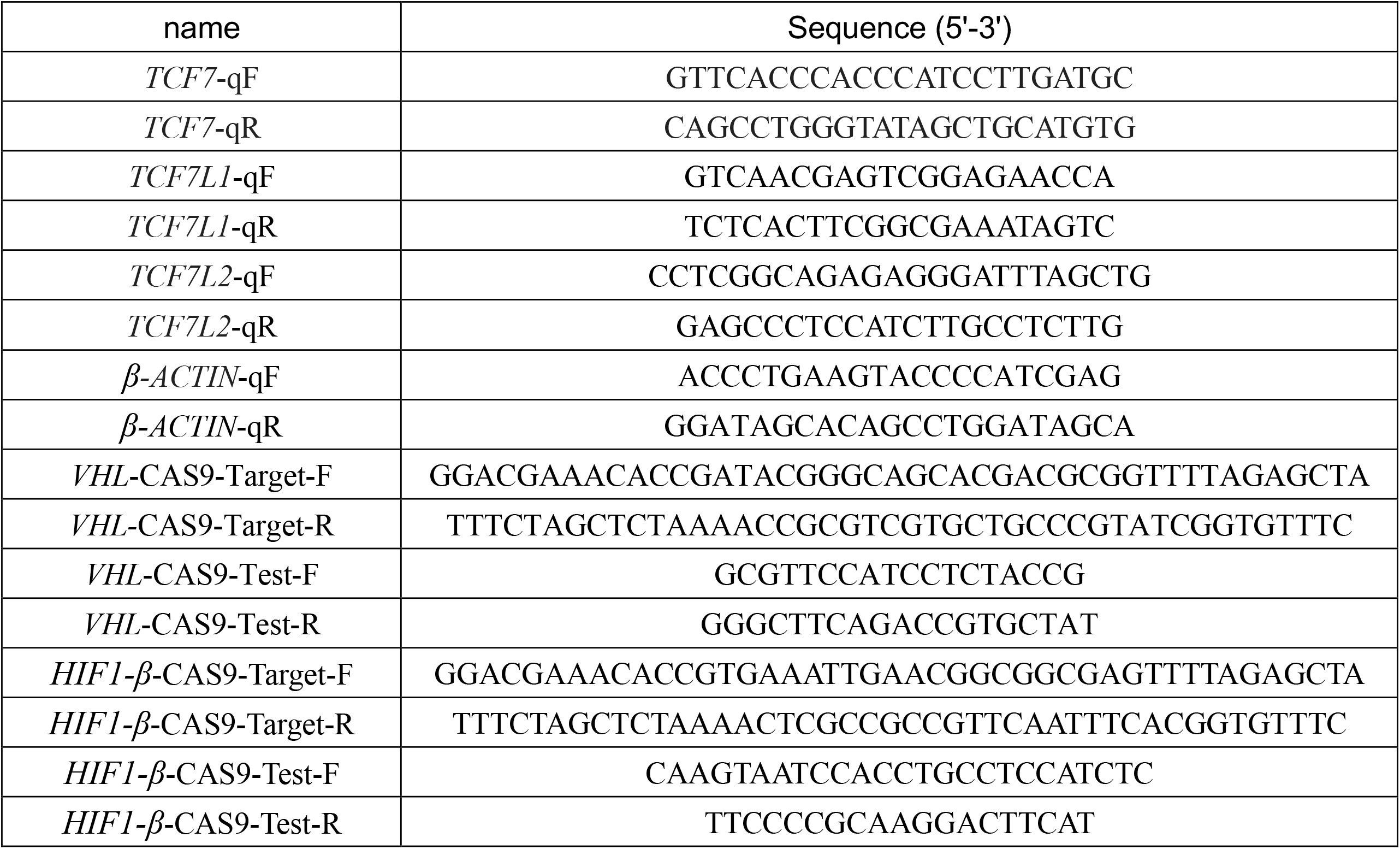
Primers and sequence information.

## Notes

### Competing Interest Statement

The authors have declared no competing interest.

## Reference

Anastas, J.N., and Moon, R.T. (2013). WNT signalling pathways as therapeutic targets in cancer. Nat Rev Cancer 13, 11–26. 10.1038/nrc3419.

Beretta, C.A., Dross, N., Bankhead, P., and Carl, M. (2013). The ventral habenulae of zebrafish develop in prosomere 2 dependent on Tcf7l2 function. Neural Dev 8, 19. 10.1186/1749-8104-8-19.

Berndt, J.D., Moon, R.T., and Major, M.B. (2009). Beta-catenin gets jaded and von Hippel-Lindau is to blame. Trends Biochem Sci 34, 101–104. 10.1016/j.tibs.2008.12.002.

Cadigan, K.M., and Waterman, M.L. (2012). TCF/LEFs and Wnt signaling in the nucleus. Cold Spring Harb Perspect Biol 4. 10.1101/cshperspect.a007906.

Carl, M., Bianco, I.H., Bajoghli, B., Aghaallaei, N., Czerny, T., and Wilson, S.W. (2007). Wnt/Axin1/beta-catenin signaling regulates asymmetric nodal activation, elaboration, and concordance of CNS asymmetries. Neuron 55, 393–405. 10.1016/j.neuron.2007.07.007.

Chitalia, V.C., Foy, R.L., Bachschmid, M.M., Zeng, L., Panchenko, M.V., Zhou, M.I., Bharti, A., Seldin, D.C., Lecker, S.H., Dominguez, I., and Cohen, H.T. (2008). Jade-1 inhibits Wnt signalling by ubiquitylating beta-catenin and mediates Wnt pathway inhibition by pVHL. Nat Cell Biol 10, 1208–1216. 10.1038/ncb1781.

Clevers, H., Loh, K.M., and Nusse, R. (2014). Stem cell signaling. An integral program for tissue renewal and regeneration: Wnt signaling and stem cell control. Science 346, 1248012. 10.1126/science.1248012.

Clevers, H., and Nusse, R. (2012). Wnt/beta-catenin signaling and disease. Cell 149, 1192–1205. 10.1016/j.cell.2012.05.012.

Cole, M.F., Johnstone, S.E., Newman, J.J., Kagey, M.H., and Young, R.A. (2008). Tcf3 is an integral component of the core regulatory circuitry of embryonic stem cells. Genes Dev 22, 746–755. 10.1101/gad.1642408.

Concha, M.L., and Wilson, S.W. (2001). Asymmetry in the epithalamus of vertebrates. J Anat 199, 63–84. 10.1046/j.1469-7580.2001.19910063.x.

Doumpas, N., Lampart, F., Robinson, M.D., Lentini, A., Nestor, C.E., Cantu, C., and Basler, K. (2019). TCF/LEF dependent and independent transcriptional regulation of Wnt/beta-catenin target genes. EMBO J 38. 10.15252/embj.201798873.

Du, J., Zhang, D., Zhang, W., Ouyang, G., Wang, J., Liu, X., Li, S., Ji, W., Liu, W., and Xiao, W. (2015). pVHL Negatively Regulates Antiviral Signaling by Targeting MAVS for Proteasomal Degradation. J Immunol 195, 1782–1790. 10.4049/jimmunol.1500588.

Du, J., Zhang, J., He, T., Li, Y., Su, Y., Tie, F., Liu, M., Harte, P.J., and Zhu, A.J. (2016). Stuxnet Facilitates the Degradation of Polycomb Protein during Development. Dev Cell 37, 507–519. 10.1016/j.devcel.2016.05.013.

Duan, D.R., Pause, A., Burgess, W.H., Aso, T., Chen, D.Y., Garrett, K.P., Conaway, R.C., Conaway, J.W., Linehan, W.M., and Klausner, R.D. (1995). Inhibition of transcription elongation by the VHL tumor suppressor protein. Science 269, 1402–1406. 10.1126/science.7660122.

Feng, Q., Zou, X., Lu, L., Li, Y., Liu, Y., Zhou, J., and Duan, C. (2012). The Stress-Response Gene redd1 Regulates Dorsoventral Patterning by Antagonizing Wnt/beta-catenin Activity in Zebrafish. Plos One 7, e52674. 10.1371/journal.pone.0052674.

Gao, C., Cao, W., Bao, L., Zuo, W., Xie, G., Cai, T., Fu, W., Zhang, J., Wu, W., Zhang, X., and Chen, Y.G. (2010). Autophagy negatively regulates Wnt signalling by promoting Dishevelled degradation. Nat Cell Biol 12, 781–790. 10.1038/ncb2082.

Giles, R.H., Lolkema, M.P., Snijckers, C.M., Belderbos, M., van der Groep, P., Mans, D.A., van Beest, M., van Noort, M., Goldschmeding, R., van Diest, P.J., et al. (2006). Interplay between VHL/HIF1alpha and Wnt/beta-catenin pathways during colorectal tumorigenesis. Oncogene 25, 3065–3070. 10.1038/sj.onc.1209330.

Gnarra, J.R., Ward, J.M., Porter, F.D., Wagner, J.R., Devor, D.E., Grinberg, A., Emmert-Buck, M.R., Westphal, H., Klausner, R.D., and Linehan, W.M. (1997). Defective placental vasculogenesis causes embryonic lethality in VHL-deficient mice. Proc Natl Acad Sci U S A 94, 9102–9107. 10.1073/pnas.94.17.9102.

Goentoro, L., and Kirschner, M.W. (2009). Evidence that fold-change, and not absolute level, of beta-catenin dictates Wnt signaling. Mol Cell 36, 872–884. 10.1016/j.molcel.2009.11.017.

Gossage, L., Eisen, T., and Maher, E.R. (2015). VHL, the story of a tumour suppressor gene. Nat Rev Cancer 15, 55–64. 10.1038/nrc3844.

Guglielmi, L., Buhler, A., Moro, E., Argenton, F., Poggi, L., and Carl, M. (2020). Temporal control of Wnt signaling is required for habenular neuron diversity and brain asymmetry. Development 147. 10.1242/dev.182865.

Guo, J., Chakraborty, A.A., Liu, P., Gan, W., Zheng, X., Inuzuka, H., Wang, B., Zhang, J., Zhang, L., Yuan, M., et al. (2016). pVHL suppresses kinase activity of Akt in a proline-hydroxylation-dependent manner. Science 353, 929–932. 10.1126/science.aad5755.

Hiyama, H., Yokoi, M., Masutani, C., Sugasawa, K., Maekawa, T., Tanaka, K., Hoeijmakers, J.H., and Hanaoka, F. (1999). Interaction of hHR23 with S5a. The ubiquitin-like domain of hHR23 mediates interaction with S5a subunit of 26 S proteasome. J Biol Chem 274, 28019–28025. 10.1074/jbc.274.39.28019.

Husken, U., and Carl, M. (2013). The Wnt/beta-catenin signaling pathway establishes neuroanatomical asymmetries and their laterality. Mech Dev 130, 330–335. 10.1016/j.mod.2012.09.002.

Husken, U., Stickney, H.L., Gestri, G., Bianco, I.H., Faro, A., Young, R.M., Roussigne, M., Hawkins, T.A., Beretta, C.A., Brinkmann, I., et al. (2014). Tcf7l2 is required for left-right asymmetric differentiation of habenular neurons. Curr Biol 24, 2217–2227. 10.1016/j.cub.2014.08.006.

Ishitani, T., Matsumoto, K., Chitnis, A.B., and Itoh, M. (2005). Nrarp functions to modulate neural-crest-cell differentiation by regulating LEF1 protein stability. Nat Cell Biol 7, 1106–1112. 10.1038/ncb1311.

Ivan, M., Kondo, K., Yang, H., Kim, W., Valiando, J., Ohh, M., Salic, A., Asara, J.M., Lane, W.S., and Kaelin, W.G., Jr. (2001). HIFalpha targeted for VHL-mediated destruction by proline hydroxylation: implications for O2 sensing. Science 292, 464–468. 10.1126/science.1059817.

Iwai, K., Yamanaka, K., Kamura, T., Minato, N., Conaway, R.C., Conaway, J.W., Klausner, R.D., and Pause, A. (1999). Identification of the von Hippel-lindau tumor-suppressor protein as part of an active E3 ubiquitin ligase complex. Proc Natl Acad Sci U S A 96, 12436–12441. 10.1073/pnas.96.22.12436.

Jaakkola, P., Mole, D.R., Tian, Y.M., Wilson, M.I., Gielbert, J., Gaskell, S.J., von Kriegsheim, A., Hebestreit, H.F., Mukherji, M., Schofield, C.J., et al. (2001). Targeting of HIF-alpha to the von Hippel-Lindau ubiquitylation complex by O2-regulated prolyl hydroxylation. Science 292, 468–472. 10.1126/science.1059796.

Kaelin, W.G. (2007). Von Hippel-Lindau disease. Annu Rev Pathol 2, 145–173. 10.1146/annurev.pathol.2.010506.092049.

Kaidi, A., Williams, A.C., and Paraskeva, C. (2007). Interaction between beta-catenin and HIF-1 promotes cellular adaptation to hypoxia. Nat Cell Biol 9, 210–217. 10.1038/ncb1534.

Kim, C.H., Oda, T., Itoh, M., Jiang, D., Artinger, K.B., Chandrasekharappa, S.C., Driever, W., and Chitnis, A.B. (2000). Repressor activity of Headless/Tcf3 is essential for vertebrate head formation. Nature 407, 913–916. 10.1038/35038097.

Kimmel, C.B., Ballard, W.W., Kimmel, S.R., Ullmann, B., and Schilling, T.F. (1995). Stages of embryonic development of the zebrafish. Dev Dyn 203, 253–310. 10.1002/aja.1002030302.

Kuan, Y.S., Roberson, S., Akitake, C.M., Fortuno, L., Gamse, J., Moens, C., and Halpern, M.E. (2015). Distinct requirements for Wntless in habenular development. Dev Biol 406, 117–128. 10.1016/j.ydbio.2015.06.006.

Li, V.S.W., Ng, S.S., Boersema, P.J., Low, T.Y., Karthaus, W.R., Gerlach, J.P., Mohammed, S., Heck, A.J.R., Maurice, M.M., Mahmoudi, T., and Clevers, H. (2012). Wnt Signaling through Inhibition of beta-Catenin Degradation in an Intact Axin1 Complex. Cell 149, 1245–1256. 10.1016/j.cell.2012.05.002.

Lin, X.Y., and Xu, X.L. (2009). Distinct functions of Wnt/beta-catenin signaling in KV development and cardiac asymmetry. Development 136, 207–217. 10.1242/dev.029561.

MacDonald, B.T., Tamai, K., and He, X. (2009). Wnt/beta-catenin signaling: components, mechanisms, and diseases. Dev Cell 17, 9–26. 10.1016/j.devcel.2009.06.016.

Mazumdar, J., O’Brien, W.T., Johnson, R.S., LaManna, J.C., Chavez, J.C., Klein, P.S., and Simon, M.C. (2010). O2 regulates stem cells through Wnt/beta-catenin signalling. Nat Cell Biol 12, 1007–1013. 10.1038/ncb2102.

Merrill, B.J., Pasolli, H.A., Polak, L., Rendl, M., Garcia-Garcia, M.J., Anderson, K.V., and Fuchs, E. (2004). Tcf3: a transcriptional regulator of axis induction in the early embryo. Development 131, 263–274. 10.1242/dev.00935.

Nusse, R., and Clevers, H. (2017). Wnt/beta-Catenin Signaling, Disease, and Emerging Therapeutic Modalities. Cell 169, 985–999. 10.1016/j.cell.2017.05.016.

Ohh, M., Yauch, R.L., Lonergan, K.M., Whaley, J.M., Stemmer-Rachamimov, A.O., Louis, D.N., Gavin, B.J., Kley, N., Kaelin, W.G., Jr., and Iliopoulos, O. (1998). The von Hippel-Lindau tumor suppressor protein is required for proper assembly of an extracellular fibronectin matrix. Mol Cell 1, 959–968. 10.1016/s1097-2765(00)80096-9.

Petersen, C.P., and Reddien, P.W. (2009). Wnt signaling and the polarity of the primary body axis. Cell 139, 1056–1068. 10.1016/j.cell.2009.11.035.

Phillips, B.T., and Kimble, J. (2009). A new look at TCF and beta-catenin through the lens of a divergent C. elegans Wnt pathway. Dev Cell 17, 27–34. 10.1016/j.devcel.2009.07.002.

Saini, S., Majid, S., and Dahiya, R. (2011). The complex roles of Wnt antagonists in RCC. Nat Rev Urol 8, 690–699. 10.1038/nrurol.2011.146.

Sakata, E., Yamaguchi, Y., Kurimoto, E., Kikuchi, J., Yokoyama, S., Yamada, S., Kawahara, H., Yokosawa, H., Hattori, N., Mizuno, Y., et al. (2003). Parkin binds the Rpn10 subunit of 26S proteasomes through its ubiquitin-like domain. EMBO Rep 4, 301–306. 10.1038/sj.embor.embor764.

Shy, B.R., Wu, C.I., Khramtsova, G.F., Zhang, J.Y., Olopade, O.I., Goss, K.H., and Merrill, B.J. (2013). Regulation of Tcf7l1 DNA binding and protein stability as principal mechanisms of Wnt/beta-catenin signaling. Cell Rep 4, 1–9. 10.1016/j.celrep.2013.06.001.

Stamos, J.L., and Weis, W.I. (2013). The beta-catenin destruction complex. Cold Spring Harb Perspect Biol 5, a007898. 10.1101/cshperspect.a007898.

Steinhart, Z., and Angers, S. (2018). Wnt signaling in development and tissue homeostasis. Development 145. 10.1242/dev.146589.

Turner, K.J., Bracewell, T.G., and Hawkins, T.A. (2014). Anatomical dissection of zebrafish brain development. Methods Mol Biol 1082, 197–214. 10.1007/978-1-62703-655-9_14.

Upadhya, S.C., and Hegde, A.N. (2003). A potential proteasome-interacting motif within the ubiquitin-like domain of parkin and other proteins. Trends Biochem Sci 28, 280–283. 10.1016/S0968-0004(03)00092-6.

van Rooijen, E., Voest, E.E., Logister, I., Korving, J., Schwerte, T., Schulte-Merker, S., Giles, R.H., and van Eeden, F.J. (2009). Zebrafish mutants in the von Hippel-Lindau tumor suppressor display a hypoxic response and recapitulate key aspects of Chuvash polycythemia. Blood 113, 6449–6460. 10.1182/blood-2008-07-167890.

Wu, D., Potluri, N., Lu, J., Kim, Y., and Rastinejad, F. (2015). Structural integration in hypoxia-inducible factors. Nature 524, 303–308. 10.1038/nature14883.

Yamada, M., Ohnishi, J., Ohkawara, B., Iemura, S., Satoh, K., Hyodo-Miura, J., Kawachi, K., Natsume, T., and Shibuya, H. (2006). NARF, an nemo-like kinase (NLK)-associated ring finger protein regulates the ubiquitylation and degradation of T cell factor/lymphoid enhancer factor (TCF/LEF). J Biol Chem 281, 20749–20760. 10.1074/jbc.M602089200.

Yi, F., Pereira, L., Hoffman, J.A., Shy, B.R., Yuen, C.M., Liu, D.R., and Merrill, B.J. (2011). Opposing effects of Tcf3 and Tcf1 control Wnt stimulation of embryonic stem cell self-renewal. Nat Cell Biol 13, 762–770. 10.1038/ncb2283.

Zhang, H., Rong, X., Wang, C., Liu, Y., Lu, L., Li, Y., Zhao, C., and Zhou, J. (2020). VBP1 modulates Wnt/beta-catenin signaling by mediating the stability of the transcription factors TCF/LEFs. J Biol Chem 295, 16826–16839. 10.1074/jbc.RA120.015282.

Zhang, J., Wu, T., Simon, J., Takada, M., Saito, R., Fan, C., Liu, X.D., Jonasch, E., Xie, L., Chen, X., et al. (2018). VHL substrate transcription factor ZHX2 as an oncogenic driver in clear cell renal cell carcinoma. Science 361, 290–295. 10.1126/science.aap8411.

Zhang, P., Yao, Q., Lu, L., Li, Y., Chen, P.J., and Duan, C. (2014). Hypoxia-inducible factor 3 is an oxygen-dependent transcription activator and regulates a distinct transcriptional response to hypoxia. Cell Rep 6, 1110–1121. 10.1016/j.celrep.2014.02.011.

